# Identification of a transcriptional signature found in multiple models of ASD and related disorders

**DOI:** 10.1101/2022.01.14.476375

**Authors:** Samuel Thudium, Katherine Palozola, Eloise L’Her, Erica Korb

## Abstract

Epigenetic regulation plays a critical role in many neurodevelopmental disorders, including Autism Spectrum Disorder (ASD). In particular, many such disorders are the result of mutations in genes that encode chromatin modifying proteins. However, while these disorders share many features, it is unclear whether they also share gene expression disruptions resulting from the aberrant regulation of chromatin. We examined 5 chromatin modifiers that are all linked to ASD despite their different roles in regulating chromatin. Specifically, we depleted Ash1L, Chd8, Crebbp, Ehmtl, and Nsd1 in parallel in a highly controlled neuronal culture system. We then identified sets of shared genes, or transcriptional signatures, that are differentially expressed following loss of multiple ASD-linked chromatin modifiers. We examined the functions of genes within the transcriptional signatures and found an enrichment in many neurotransmitter transport genes and activity-dependent genes. In addition, these genes are enriched for specific chromatin features such as bivalent domains that allow for highly dynamic regulation of gene expression. The downregulated transcriptional signature is also observed within multiple mouse models of neurodevelopmental disorders that result in ASD, but not those only associated with intellectual disability. Finally, the downregulated transcriptional signature can distinguish between neurons generated from iPSCs derived from healthy donors and idiopathic ASD patients through RNA-deconvolution, demonstrating that this gene set is relevant to the human disorder. This work identifies a transcriptional signature that is found within many neurodevelopmental syndromes, helping to elucidate the link between epigenetic regulation and the underlying cellular mechanisms that result in ASD.

## Introduction

Neurodevelopmental Disorders (NDDs) that result in Autism Spectrum Disorder (ASD) are caused by both environmental and genetic factors. Even within the subset of disorders that have a clear genetic cause, each individual syndrome stems from a unique mutation in an increasingly long list of ASD susceptibility genes. Such heterogeneity has made it difficult to develop a unifying model of the disruptions that lead to shared phenotypes or to develop treatments that address shared underlying causes. Interestingly, recent studies demonstrated that a disproportionate number of ASD susceptibility genes encode epigenetic regulators (Iossifov et al., 2014; O’Roak et al., 2012; Parikshak et al., 2013; De Rubeis et al., 2014). In particular, many such mutations are found in proteins that regulate chromatin, the complex of DNA and histone proteins that helps to regulate transcription.

Histones are regulated by numerous posttranslational modifications such as acetylation, methylation, and many others, which ultimately affect transcription. These modifications recruit transcriptional regulators and allow chromatin to transition between the open and closed states that are permissive or repressive to transcription, thus providing a complex code that regulates gene expression (Berger, 2007; Jenuwein and Allis, 2001; Strahl and Allis, 2000; Turner, 2000). The importance of this ‘histone code’ or ‘language’ is becoming increasingly appreciated in neuroscience, from its function in memory formation to its involvement in neurodevelopmental disorders (Borrelli et al., 2008; Peixoto and Abel, 2013; Rangasamy et al., 2013). However, it remains unclear if the different forms of syndromic ASD that result from mutations in distinct chromatin regulators share transcriptional disruptions.

Determining whether disruption of multiple syndromic ASD-linked chromatin modifiers with disparate functions leads to overlapping gene expression changes presents multiple challenges. Thus far, such chromatin modifiers have been analyzed individually in different systems, but never in parallel in a controlled genetic background. As a result, while our understanding of these disorders has improved drastically in recent years, previous studies were not designed to allow for a comparison between different causes of ASD or identify common pathways that underlie shared phenotypes. Work examining the effects of loss of these chromatin modifying proteins in animals is also confounded by the full body and lifelong loss of these proteins throughout development. Thus, the complexity of the compensatory response and other related health effects may occlude any relevant transcriptional signature that could answer these outstanding questions. Finally, many neurodevelopmental disorders result in a range of phenotypes and often cause *both* ASD and intellectual disability (ID), so identifying which underlying epigenetic disruptions are associated with one or both phenotypes poses additional hurdles.

To overcome these challenges, we developed a primary neuronal culture system that allows us to study multiple syndromic ASD-linked chromatin modifiers in parallel, in a controlled genetic background, with the goal of defining the common gene expression patterns caused by disruption of these proteins. Remarkably, we found that loss of five such proteins, despite having a diverse array of functions, all cause disruption of similar sets of genes, particularly for downregulated genes. We termed the sets of genes disrupted by depletion of the majority of the chromatin modifiers *transcriptional signatures.* These signatures encoded genes relevant to synaptic function, including activity-dependent genes. Further, these genes shared several chromatin features, including bivalent domains, that may make them particularly sensitive to chromatin disruptions. In addition, the downregulated gene signature that we identified in cultured neurons is also present in animal models of NDDs that result in ASD. However, they are not present in disorders that result in ID in the absence of ASD, or in a neurodegenerative disorder. Finally, we mapped this signature onto gene expression data from human induced pluripotent stem cell (iPSC) derived neurons and found that it is able to distinguish between control and idiopathic ASD patient cells. These data indicate that common sets of genes are disrupted after loss of multiple chromatin-associated proteins linked to ASD. Moreover, these genes control critical neuronal functions, share chromatin features, and are also disrupted in multiple mouse models and in ASD iPSC derived neurons. Defining this signature provides novel insights into the cellular disruptions that contribute to ASD and how mutations in a diverse array of histone-modifying enzymes can lead to common phenotypic outputs.

## Results

### Defining gene expression profiles of syndromic ASD-linked chromatin modifiers

We sought to determine if the loss of different chromatin modifying enzymes linked to related neurodevelopmental disorders results in a common transcriptional signature. To define such a signature, we focused on five chromatin modifiers, Ash1L, Chd8, Crebbp, Ehmt1, and Nsd1, associated with syndromes that include intellectual disability and ASD traits. (**Table 1**). Of the many chromatin regulators linked to such disorders (Neale et al., 2012; O’Roak et al., 2012; Sanders et al., 2012; De Rubeis et al., 2014; Iossifov et al., 2014), we chose the subset that led to well-defined syndromes caused by loss-of-function mutations or deletions (Abrahams et al., 2013). This ensured we examined proteins whose loss results in NDDs with high penetrance. Further, these 5 proteins among the top ASD risk genes as identified by TADA analysis (Fu et al., 2021) with 4 of the 5 in the top 100 susceptibility genes for idiopathic ASD and in the same gene expression module (based on BrainSpan data) which contains the greatest enrichment of ASD susceptibility genes (Ji et al., 2016). We further selected for chromatin regulators with mouse models that recapitulate the features of the associated disorder to ensure that mouse neuronal models are an appropriate system in which to study their function (Benevento et al., 2016; Bernier et al., 2014; Coupry et al., 2002; Eram et al., 2015; Gao et al., 2021; Kleefstra et al., 2006, 2009; Kurotaki et al., 2002; Neale et al., 2012; Niikawa, 2004; Shen et al., 2019). Finally, we selected a set of chromatin modifiers that have a diverse array of functions in chromatin. These proteins target different substrates in chromatin including modifying different histone proteins and residues. They also perform a diverse array of functions such adding different types of modifications. For example, CREBBP acetylates histones, Emht1 methylates H3K9, while Ash1L and Nsd1 promote different methylation states of H3K36 (Jin et al., 2011; Miyazaki et al., 2013; Qiao et al., 2011; Tachibana et al., 2008; Thompson et al., 2008). Finally, some are associated with active gene expression (Crebbp, Ash1l, Nsd1) while others are associated with repressive gene expression (Ehmt1, Chd8). Thus, we would not expect these proteins to target the same set of genes or their loss to result in similar transcriptional profiles solely based on having shared functions in modulating chromatin. Instead, the major commonality between these proteins is that their disruption leads to ID and ASD, so any overlapping gene expression changes are more likely to be relevant to shared phenotypic output.

**Table 1.** Candidate proteins. Functions and associated disorders of 5 chromatin modifiers chosen for analysis. All proteins have distinct functions in regulating histones. Mutation or deletion of all candidates results in well-defined neurodevelopmental syndromes that include features such as intellectual disability (ID) and Autism Spectrum Disorder (ASD). Mouse (Ms) column indicates a mouse models show expected phenotypes.

| Gene ID | Function | Target (mark) | Associated Syndrome | ASD | ID | Ms |
| --- | --- | --- | --- | --- | --- | --- |
| *Ash1L* | Histone lysine methyltransferase | H3K36 (me3) | Emerging MCA/ID Disorder | X | X | X |
| *Chd8* | Chromodomain helicase | Chromatin remodeler; recruitment of other complexes | Autism 18; Charge Syndrome | X | X | X |
| *Crebbp* | Histone acetyltransferase | Promiscuous H3 lysines (Ac) | Rubinstein-Taybi Syndrome | X | X | X |
| *Ehmt1* | Histone lysine methyltransferase | H3K9 (me1/me2) | Kleefstra Syndrome | X | X | X |
| *Nsd1* | Histone lysine methyltransferase | H3K36 (me2) | Soto Syndrome; Beckwith-Wiedemann Syndrome | X | X | X |

To study the effects of the loss of these chromatin modifiers in parallel, we used a primary neuronal culture system and lentiviral shRNA knockdown of each chromatin modifier (**Fig. 1A**). There are several advantages to this approach: 1) This culture method generates a purely neuronal population (Korb et al., 2015) and thus avoids the heterogeneity of brain tissue and the compounding effects of a system-wide knockout; 2) Neurons are cultured from embryonic mouse brains which allows for the investigation of early neuronal development time points that are relevant to the onset of NDDs and ASD; 3) Neurons are analyzed 5 days after knockdown thus avoiding long-term compensatory responses resulting from life-long loss of function; 4) For each biological replicate, each candidate gene is knocked down simultaneously from neurons cultured from the same animal, thereby controlling for both genetic background, developmental time point, and variation between animals; and 5) Multiple replicates can be generated and processed in parallel to allow for a high degree of rigor using true biological replicates (each coming from separate liters of mice) while also minimizing technical variability. While ID and ASD can be caused by atypical brain region connectivity that cannot be detected in our system, our goal is to define the underlying cellular mechanisms *within* neurons that ultimately lead to wider disruptions.

**Figure 1.**
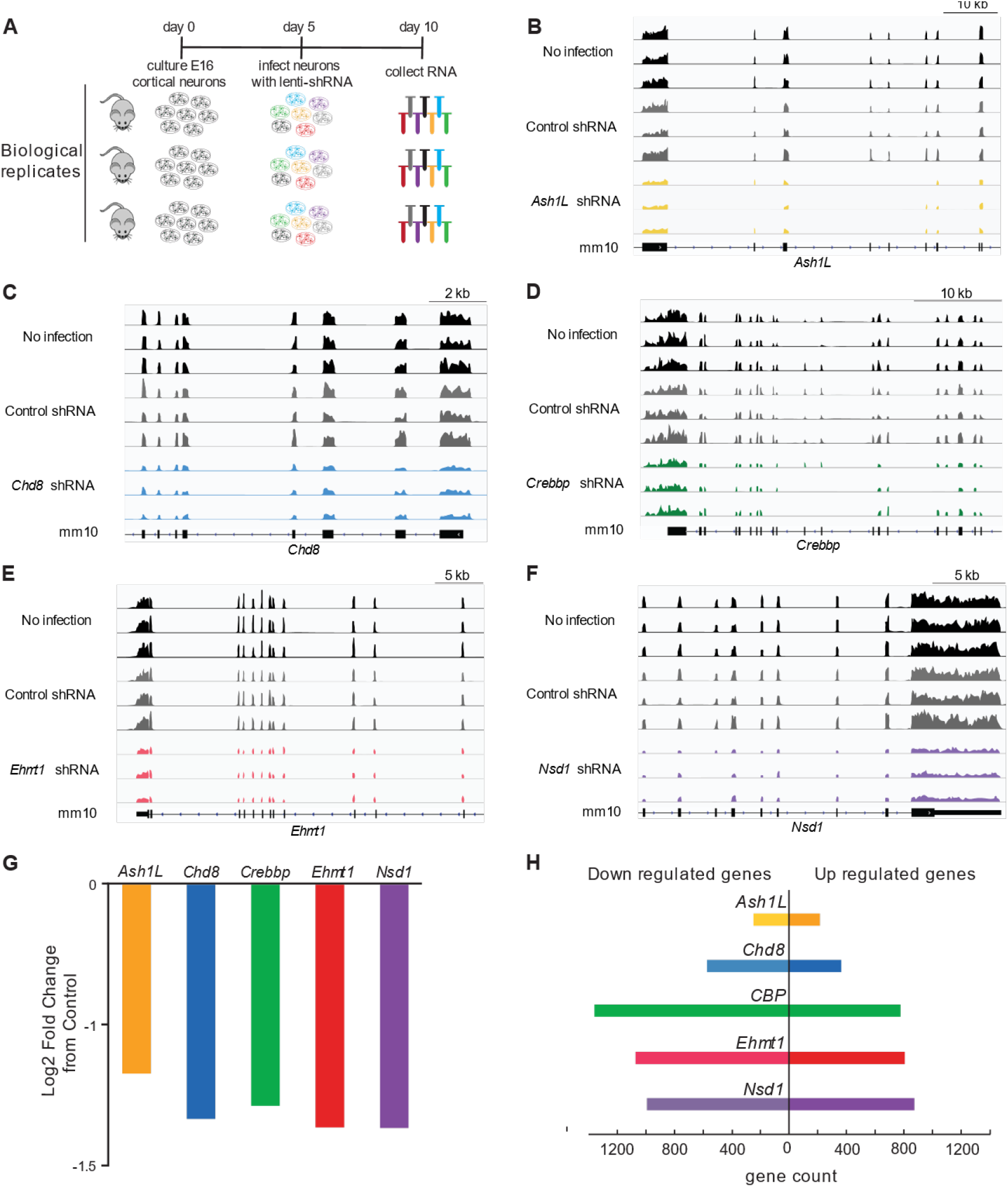
System for comparison of gene expression profiles of chromatin modifiers. (A) Primary neuronal culture system used to analyze gene expression changes after knockdown of ASD-linked chromatin modifiers. (B-F) Gene tracks of chromatin modifier genes after knockdown. (G) Fold change in transcript expression of targets from RNA-seq data. (H) Number of genes down and upregulated by knockdown of 5 ASD-linked chromatin-associated proteins. N=3 replicates.

Primary cultured neurons were infected with lentiviruses containing shRNAs targeting each syndromic ASD-linked chromatin modifier or with a non-targeting control shRNA at five days in culture. Neurons were then collected 5 days after infection to allow for robust depletion of target proteins. We used RT-qPCR to examine the degree of knockdown achieved through lentiviral infection and confirmed depletion of all target transcripts (**Supplemental Fig. 1A**). We further confirmed knockdown of the target proteins in all cases where antibodies were available (**Supplemental Fig. 1B**). We then used RNA-sequencing (RNA-seq) to define gene expression changes resulting from knockdown of all five chromatin-associated proteins and further confirmed knockdown through sequencing data (**Fig. 1B-G**). Having ensured robust knockdown through multiple approaches, we analyzed all gene expression after depletion of each chromatin modifier. As expected, in all cases knockdown resulted in robust gene expression changes, with genes both increased and decreased in expression compared to non-targeting control shRNA infection (**Fig. 1H, Supplemental Figure 2**).

### Overlap of gene expression changes caused by loss of ASD-linked chromatin modifiers

To examine potential overlap in the resulting gene expression changes, we took several approaches. First, we examined the direct overlap of each possible pairwise comparison and used a hypergeometric test to determine the significance of the number of overlapping genes. Remarkably, we found that direct comparison of every downregulated gene set was significant and the same was true of each upregulated gene set overlap (**Fig. 2A**). Conversely, comparison of each upregulated gene set with each downregulated gene set yielded almost no significant overlaps. These findings indicate that many of the same genes are differentially regulated by these 5 ASD-linked chromatin modifiers. This is particularly remarkable because these 5 chromatin modifiers were chosen specifically based on their divergent functions in regulating chromatin and thus their depletion would not necessarily be expected to have similar effects on gene expression. Instead, the significance of these overlaps indicates that this subset of genes may be particularly sensitive to disruption of ASD-linked chromatin modifiers in neurons.

**Figure 2.**
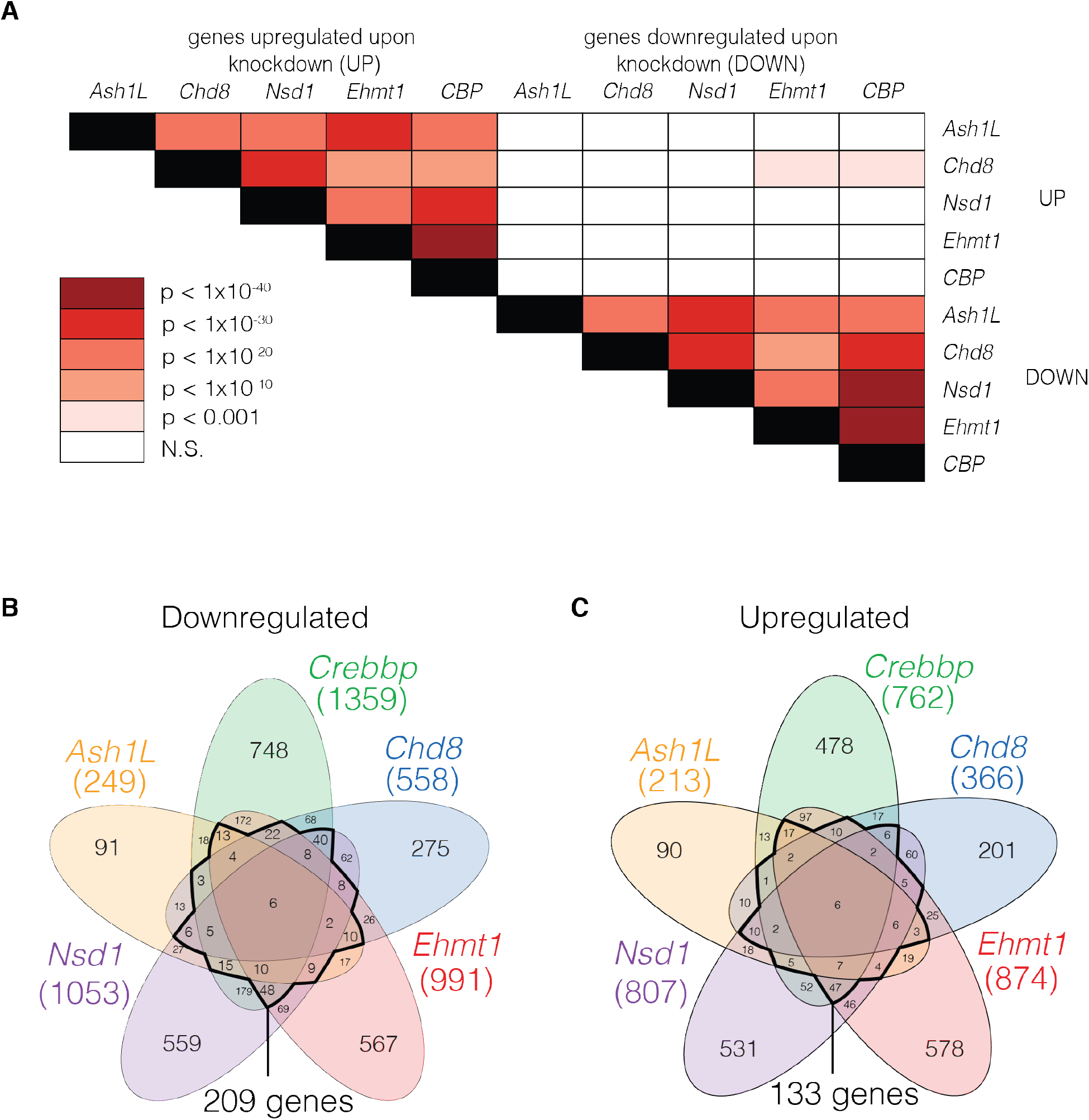
Overlap of genes that are differentially expressed following knockdown of ASD-linked chromatin modifiers. (A) Significance of overlap of down and upregulated genes after knockdown. Heatmap indicates significance level by hypergeometric testing. (B-C) Overlap of genes that are downregulated (B) or upregulated (C) by multiple targets. Dark line indicates subset of genes differentially expressed in at least 3 of 5 gene sets. Overlap significance based on hypergeometric tests.

We next examined all higher-order intersections between these gene sets to determine which genes are commonly disrupted in response to knockdown of multiple ID/ASD-linked chromatin modifiers (**Fig. 2B-C**). Only 6 genes were common between all datasets for both up (Trak2, Snap29, Prickle1, Ccnt1, Ppig, and Fam214b) and downregulated (Fcgrt, Dbp, AI464131, Creb3l1, Ret, and Isoc1) genes. However, 209 downregulated and 133 upregulated genes were shared by knockdown of the majority (at least 3 of the 5) of the ASD-linked chromatin modifiers. Thus, while these ASD-linked chromatin modifiers all have different functions in regulating chromatin and target different histone residues and genomic regions, the gene expression changes resulting from their depletion converge on common subsets of genes, particularly for downregulated genes. For simplicity, we will refer to the sets of genes that are up or downregulated in response to the majority of these chromatin modifiers as *transcriptional signatures*. Given the greater number of downregulated genes shared between the 5 chromatin modifiers analyzed, we will largely focus on the downregulated gene signature here and include all analyses of the upregulated signature in supplemental figures.

### Functional enrichments in the transcriptional signatures

We first sought to define the gene functions encoded by the up and down transcriptional signatures using Gene Ontology (GO) analysis. We found that the top enriched terms for the downregulated signature included processes such as neurotransmitter transport and nervous system development, as would be expected for a gene set that is linked to ASD and ID (**Fig. 3A**). To ensure that the functional groups we identified weren’t unique to one type of analysis, we also performed GeneWalk (Ietswaart et al., 2021) which generates a gene regulatory and GO-term network comprising all input genes. Significant functions are determined based on the similarity of vector representations for each gene to identify gene functions that are relevant to the biological context of a given experiment. We then used REVIGO (Supek et al., 2011) to remove redundant outputs and cluster related functions, and labeled each resulting cluster with an identifier that encompassed the GO terms included (**Fig. 3B**). The GeneWalk output fit remarkably well with the standard GO analysis, demonstrating that these functional groups are enriched in the downregulated transcriptional signature regardless of the methods used.

**Figure 3.**
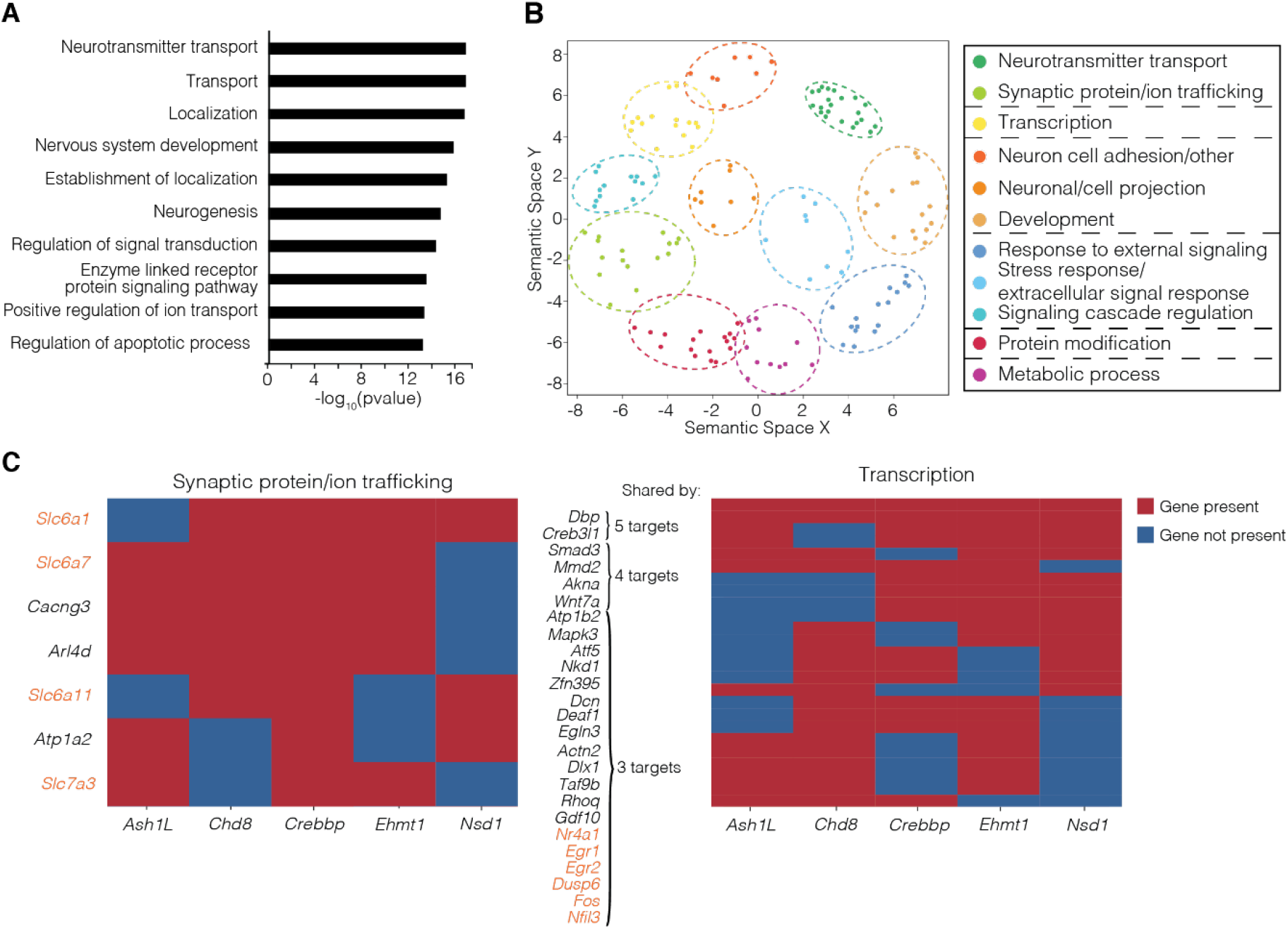
Function of downregulated transcriptional signature genes. (A) Gene Ontology analysis of downregulated transcriptional signature gene function. (B) GeneWalk analysis followed by Revigo clustering of downregulated transcriptional signature genes. (C) Genes contributing to main gene ontology clusters that are differentially expressed after knockdown of 3 or more ASD-linked chromatin modifiers. Solute-carrier family and activity dependent genes are shown in orange.

We next sought to define the specific genes within the transcriptional signature responsible for driving these functional enrichments. We identified the genes driving the enrichment of top GO terms and determined how many of the 5 gene sets contained these gene drivers (**Fig. 3C, Supplemental Fig. 3A**). Multiple members of the solute carrier family of genes, which regulate neurotransmitter transport, were present in at least 3 of the 5 gene sets contributing to the neurotransmitter and synaptic protein trafficking gene ontology groups. We also found that many of the genes detected within at least 3 datasets for the ‘Transcription’ and ‘Response to extracellular signal’ gene ontology groups identified through GeneWalk are well-established activity-dependent genes (also found in corresponding standard GO terms ‘regulation of signal transduction’ and ‘Regulation of apoptotic process’). To directly examine whether activity-dependent genes were significantly enriched within the downregulated transcriptional signature, we utilized a previously generated set of genes upregulated in neurons upon simulation by Brain Derived Neurotrophic Factor (BDNF) for 10 minutes (Korb et al., 2015). Upon comparing these lists, we found the downregulated transcriptional signature was significantly enriched for these genes, indicating that activity-dependent genes are among those preferentially disrupted by loss of ID/ASD-linked chromatin modifiers (**Supplemental Fig. 3B**). We then confirmed these findings in an *in vivo* system by examining gene expression changes following a recall event in activated neurons using the TRAP2 mouse model (Chen et al., 2020) (**Supplemental Fig. 3C**) Finally, we used RT-qPCR to confirm downregulation of genes of interest following knockdown of the 5 ASD associated chromatin modifiers, including several activity-dependent genes *(Fos* and *Nfil3),* a solute carrier family gene *(Slc7a3),* and a calcium channel gene *(Cacng1a)* (**Supplemental Fig. 3D**). We found that each of these genes was depleted in response to knockdown of at least 4 ASD-linked chromatin modifiers.

We performed similar analyses to identify functional groups enriched within the upregulated transcriptional signature. Using standard GO, we found that the top 10 functional groups within this signature largely included functions relevant to cell division (**Supplemental Fig. 3E**). While neurons are postmitotic, they use many cell cycle genes for regulation of neuronal maturation and migration (Frank and Tsai, 2009; Huang et al., 2010; Lim and Kaldis, 2013; Ohnuma and Harris, 2003). GeneWalk and REVIGO clustering revealed enriched clusters including neuronal maturation that corresponded to the cell division gene sets found by GO (**Supplemental Fig. 3F**).

Rather than simply starting with the transcriptional signatures defined above, we also examined the common functionally enriched groups shared between the ID/ASD-linked chromatin modifiers using a converse approach. We first performed GeneWalk on each of the 5 chromatin modifiers’ up or down differentially expressed gene sets individually, then overlapped the resulting GO terms to define a set of ontology terms common to all gene sets (**Supplemental Fig. 4A)**. REVIGO was used to cluster output terms and we identified each resulting cluster based on the GO terms included. We found functional groups for down regulated genes through this approach that were equivalent to those found through GO and GeneWalk analysis of the ASD downregulated signature, including responses to external signaling (containing activity-dependent genes) and synaptic protein trafficking (containing neurotransmitter transport genes) (**Supplemental Fig. 4B**). Similarly, by this alternate approach, we found many analogous functional clusters present in the gene ontology terms shared by all up regulated gene lists, such as cell morphology and development, (**Supplemental Fig. 4C**). Thus, multiple functional analyses indicate that gene expression changes in response to loss of ID/ASD-linked chromatin modifiers affect critical neuronal regulatory processes such as neuronal development, synaptic trafficking, and activity-dependent gene regulation.

### Chromatin features of the transcriptional signature

Having defined the functional relevance of the transcriptional signatures, we next sought to understand the features that make these genes particularly susceptible to disruption in response to knockdown of ASD-linked chromatin modifiers. We first examined their expression within control conditions and found, as expected, that they were all expressed within neurons, but otherwise include a wide-range of relative expression levels with no notable enrichment for low or high expressed genes (**Supplemental Fig. 5A-B**). Previous work has identified specific chromatin features that are shared between genes that are disrupted in neurodevelopmental disorders (Zhao et al., 2018). We therefore examined the chromatin features found within these transcriptional signatures. We used ChromHMM (Ernst and Kellis, 2017) to define the chromatin states enriched at the promoters (**Fig. 4A**) and gene bodies (**Fig. 4B**) of the downregulated transcriptional signature. We compared transcriptional signature genes with the entire genome, the genes expressed in neurons based on RNA-seq data, and all genes. As expected, the full genome was depleted of defined chromatin states relative to all gene sets examined. Interestingly, several features are distinct between the promoters of the transcriptional signature genes compared to both the promoters of all genes and the promoters of the subset of genes expressed within neurons. Mostly strikingly, promoters of the transcriptional signature genes were enriched for a bivalent state. Bivalency refers to the cooccurrence of histone modifications associated with opposing functions, and is typically defined by the presence of H3K4me3, which is associated with active gene promoters, and H3K27me3, which is associated with transcriptional repression. The synchronous presence of functionally opposing histone modifications allows for genes to be maintained in a poised state and rapidly activated in response to external signals (Bernstein et al., 2006; Voigt et al., 2013). In addition, the downregulated transcriptional signature was enriched for chromatin state corresponding to strong promoter-proximal enhancers, further indicating the presence of chromatin regulatory features that allow for the robust and highly regulated activation of target genes.

**Figure 4.**
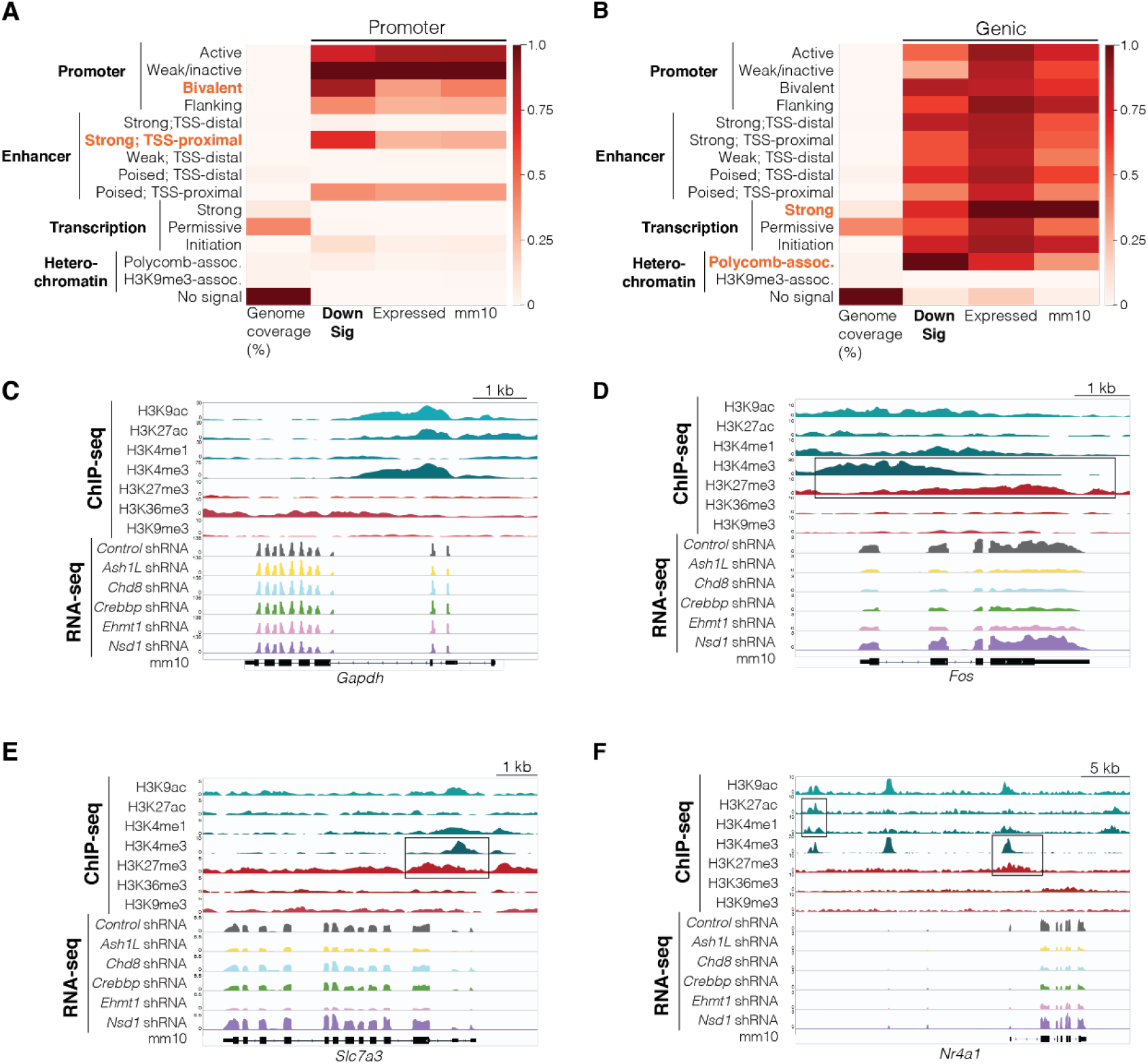
Chromatin states in transcriptional signature genes. (A) ChromHMM analysis of promoter (500 base pairs upstream of transcription start site) of downregulated transcriptional signature genes. (B) ChromHMM analysis of genic regions of transcriptional signature genes. (C) Gene track of a control gene, *Gapdh,* that is not regulated by ASD-linked chromatin modifiers. (D-F) Gene tracks of downregulated transcriptional signature genes *Fos* (D), *Slc7a3* (E), and *Nr4a1* (F) that have bivalent domains (high H3K4me3 and high H3K27me3), low H3K36me3, and strong proximal enhancer site (H3K4me1 and H3K27ac peaks upstream of *Nr4a1)* typical of downregulated transcriptional signature genes. Boxes highlight these chromatin states. TSS indicates transcription start site. Expressed indicates genes expressed in neuronal culture system. Displayed heatmaps represent overlap enrichment output values range-normalized by column.

To ensure that the enrichment of these chromatin features is not dependent on the size of the region surrounding the TSS used to define the promoter and that the analysis did not inappropriately exclude features present in broader regions, we repeated ChromHMM with an expanded region upstream of the TSS and again found similar enrichment of these chromatin states (**Supplemental Fig. 5C**). We further sought to confirm that there is an enrichment of bivalent genes within this gene signature. We used a previously defined (Court and Arnaud, 2017) set of bivalent genes and identified the subset of these that are expressed in the neuronal culture model used here. We found a highly significant overlap between these bivalent genes and the downregulated transcriptional signature (**Supplemental Fig. 5D**), confirming an enrichment of bivalent chromatin states within the genes disrupted by depletion of ASD-linked chromatin modifiers.

Next, we examined the chromatin states in genic regions of the downregulated transcriptional signature as above (**Fig. 4B**). We found an enrichment in polycomb-associated chromatin, reflective of the H3K27me3 present in gene bodies of bivalent genes. We also observed a decrease in the chromatin state associated with strong transcription, corresponding to the histone modification H3K36me3. This modification has multiple functions including promoting histone deacetylation to prevent run-away transcription and mediating splicing in neurons (Wagner and Carpenter, 2012; Xu et al., 2021). We also examined specific gene tracks of genes driving the functional gene ontology enrichments (**Fig. 4C-D**), specifically activity-dependent genes and genes regulating neurotransmitter transport that were present in the downregulated transcriptional signature. These genes contained bivalent domains with high H3K4me3 and H3K27me3 relative to ubiquitously expressed genes such as *Gapdh* (**Fig. 4C-F**). Interestingly, these genes also had low H3K36me3, and in genes containing proximal enhancers, such as *Nr4a1*, strong enhancer domains marked by H3K4me1 and H3K27ac.

We also examined chromatin states of upregulated transcriptional signature genes. Interestingly, for genic regions, many chromatin states associated with enhancers were lower in this gene set compared to all genes expressed within neurons **(Supplemental Fig. 5E**). While the down regulated signature promoter coordinates were highly enriched for bivalent domains, we found that the opposite was true at the promoters of the up regulated gene signature; this was especially apparent in the more restrictive promoter region (**Supplemental Fig. 5F).** Further, we saw robust enrichment of the active promoter state, marked by strong signals of H3K4me3, H3K9ac, and H3K27ac, in both the narrow and expanded promoter regions of the up regulated signature genes (**Supplemental Fig. 5F-G)**. Together, these findings suggest that genes found within the transcriptional signatures may be particularly susceptible to disruption due to distinct chromatin features. Downregulated genes in particular have features of a poised chromatin state, such as bivalent modifications at the promoter and modifications that support strong proximal enhancer function, while upregulated genes have modifications that confer strong promoter activity.

### Identification of the transcriptional signature in mouse models of syndromic ASD

While the neuronal cell culture model used here to define transcriptional signatures provides a highly controlled system, its relevance to ASD may not translate in the physiological context of the brain. Therefore, to determine if the transcriptional signatures defined through a primary neuronal culture model are indicative of gene expression changes occurring within the brain, we compared these signatures to gene expression changes in multiple mouse models of NDDs. We first examined a mouse model of Rubenstein-Taybi Syndrome, which is characterized by short stature, moderate to severe ID, features of ASD, and additional abnormalities such as heart and kidney defects. It is most often caused by mutations in either of the *KAT3A* genes, *Crebbp* or *p300,* which have overlapping functions as histone acetyltransferases. *Crebbp* was one of 5 chromatin modifiers used to generate the transcriptional signatures here and thus we would hypothesize that these signatures would be present in this model if they are relevant to an *in vivo* system. We used RNA-seq of cortical tissue from a mouse model containing a double knockout of the *Kat3a* genes *(Crebbp* and *p300)* (Lipinski et al., 2020) and examined the direct overlap of differentially expressed genes to the transcriptional signatures. We found a highly significant overlap between genes downregulated in the Rubenstein-Taybi mouse model and the downregulated transcriptional signature (69 genes overlap, p = 1×10^-6^) (**Fig. 5A**). As an alternate approach, we used GSEA to map the downregulated transcriptional signature genes onto a log2 fold change ranked list of all gene expression changes in *KAT3A* knockout mice. Again, we found a highly significant enrichment through this analysis with downregulated transcriptional signature genes present within highly downregulated genes in the *KAT3A* knockout mouse (**Fig. 5B**). These analyses indicate that the transcriptional signature identified in cultured mouse neurons is also detected in related animal models.

**Figure 5.**
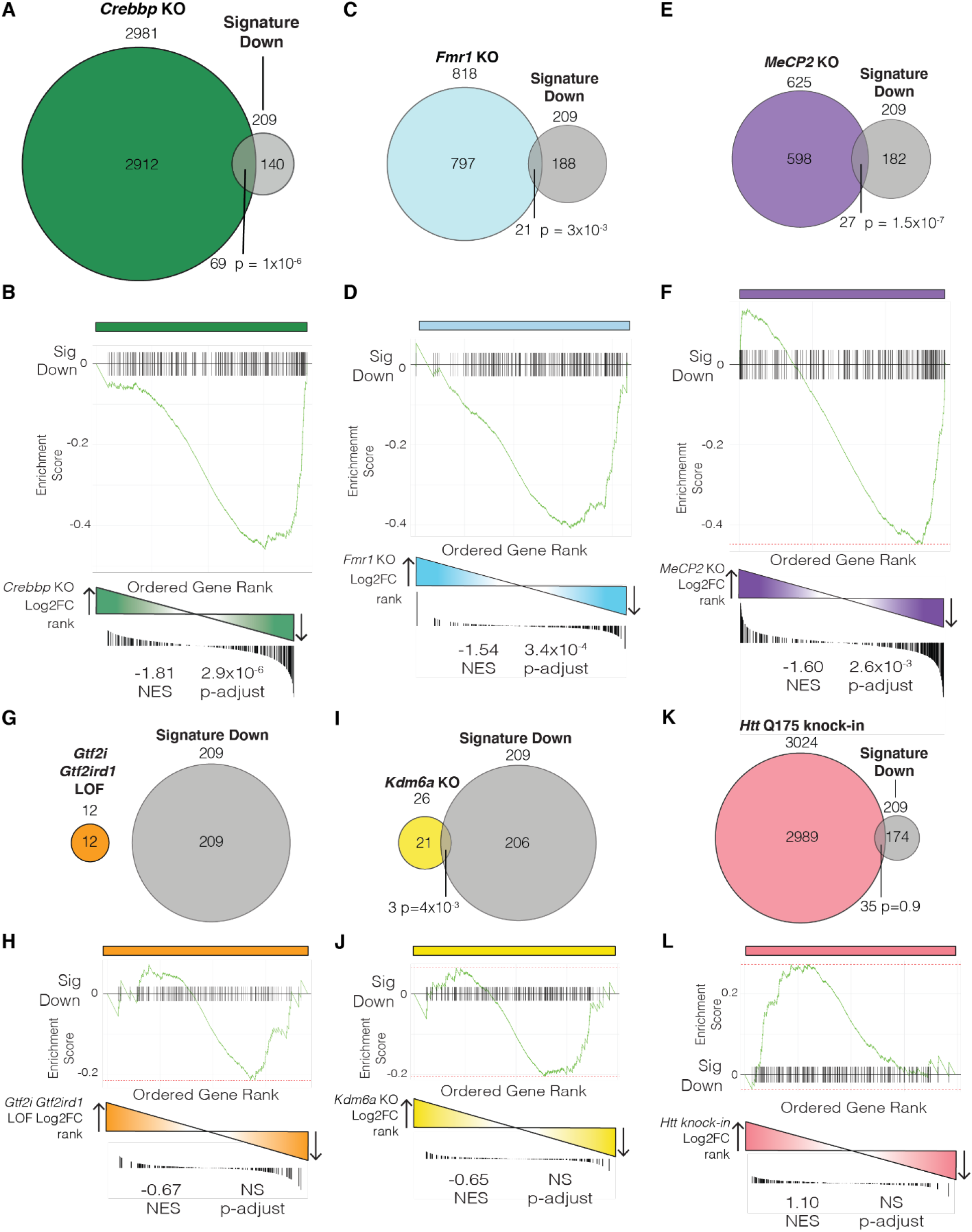
Identification of transcriptional signature in mouse models of ASD. (A-B) Overlap (A) and GSEA (B) analysis of downregulated transcriptional signature compared to differentially expressed genes in a *Kat3a* double KO mouse model of Rubinstein-Taybi Syndrome. (C-D) Overlap (C) and GSEA (D) analysis of downregulated transcriptional signature compared to differentially expressed genes in a *Fmr1* KO mouse model of FXS. (E-F). Overlap (E) and GSEA (F) analysis of downregulated transcriptional signature compared to differentially expressed genes in a *MeCP2* KO mouse model of Rett Syndrome. Overlap (G) and GSEA (H) analysis of downregulated transcriptional signature compared to differentially expressed genes in a *Gtf2i* and *Gtf2ird* double LOF mouse model of Williams Syndrome. (I-J) Overlap (I) and GSEA (J) analysis of downregulated transcriptional signature compared to differentially expressed genes in a *Kdm6a* KO mouse model of Kabuki Syndrome. (K-L) Overlap (K) and GSEA (L) analysis of downregulated transcriptional signature compared to differentially expressed genes in a *HTT* Q175 repeat knock-in mouse model of Huntington’s Disease. Overlap significance based on hypergeometric tests. NES indicates normalized enrichment score. LOF indicates loss of function.

We next examined whether the transcriptional signature is found in disorders that include ASD features but that are not directly caused by loss of the 5 chromatin modifiers we examined here. We focused on Fragile X Syndrome (FXS), a leading genetic cause of both ID and ASD. FXS is typically caused by a repeat expansion that results in loss of expression of the *Fmr1* gene that encodes the FMRP protein. FMRP has multiple functions including regulating translation of target mRNAs which encode synaptic proteins (Darnell et al., 2011; Niere et al., 2012), as well as chromatin modifiers (Korb et al., 2017), both of which are disrupted in response to loss of FMRP. We used a mouse model of FXS, containing a knockout of the *Fmr1* gene, that recapitulates many aspects of the human disorder (Korb et al., 2017; Spencer et al., 2005, 2008). We found that both by direct overlap (**Fig. 5C**) and by GSEA analysis (**Fig. 5D**), the downregulated transcriptional signature genes were significantly enriched in genes downregulated in FXS mouse cortices (21 genes overlap, p = 3×10^-3^).

Lastly, we examined a mouse model of Rett Syndrome, which results in global deficits including loss of speech, movement disruptions, and autistic features. It is typically caused by mutations in the gene encoding MECP2 which binds methylated DNA and recruits protein complexes to regulate gene expression (Good et al., 2021). We examined RNA-seq data obtained from several mouse models of Rett Syndrome, including a full knockout of MECP2, a model containing a common MECP2 patient mutation, and a heterozygous deletion of MECP2 (Jiang et al., 2021; Pacheco et al., 2017). In all cases, we detected a significant overlap of differentially downregulated genes with the downregulated transcriptional signature (27 genes overlap, p = 1.5×10^-7^ in the full KO; 22 genes overlap, p = 8.2×10^-6^ in the patient mutation model; 27 genes overlap, p = 2.2×10^-9^ in the heterozygous model) (**Fig. 5E, Supplemental Fig. 6A, C)** and a significant enrichment of the downregulated transcriptional signature by GSEA (**Fig. 5F, Supplemental Fig. 6B**). Genes downregulated in the heterozygous model were not nearly as enriched for the down regulated signature as the full knockout by GSEA (**Supplemental Fig. 6D**). Together, these data demonstrate that downregulated transcriptional signature genes are disrupted in multiple animal models of ASD, even beyond those directly related to the chromatin modifiers used to define this signature in neuronal cultures.

Given these robust findings, we asked whether the transcriptional signature is detected in any animal model of an NDD in which transcription is disrupted in neurons, or if this signature is more closely associated with specific phenotypic outcomes such as ID or ASD. To this end, we examined mouse models of Williams Syndrome, a neurodevelopmental disorder caused by deletion of a region on chromosome 7 that encompasses 26 to 28 genes. Williams syndrome results in ID but, rather than causing ASD, patients have a hypersociability phenotype caused by deletion of transcription factors *Gtf2i* and *Gtf2ird1*. Loss of these genes leads to both ID and hypersociability in animal models (Barak et al., 2019; Dai et al., 2009; Kopp et al., 2019; Segura-Puimedon et al., 2014; Young et al., 2008). Using RNA-seq data from hippocampus of a *Gtf2i Gtf2irdl* double knockout mouse model (Kopp et al., 2019), we found no overlapping gene expression changes with the downregulated transcriptional signature (**Fig. 5G)**. To confirm that this was not solely due to the small number of differentially expressed genes, we performed preranked GSEA based on log2 fold change, which considers all gene expression changes and thus is not constrained by gene list size, and again found no enrichment (**Fig. H**). As another control to ensure that any lack of overlap was not just due to the specific animal model of Williams Syndrome chosen, we repeated this analysis with the complete Williams Syndrome chromosomal deletion. We again found no significant overlap with any differentially expressed genes, and no enrichment through GSEA (**Supplemental Fig. 6E-F**).

To determine whether the lack of enrichment of the transcriptional signature in Williams Syndrome models was specific to this syndrome, we also examined a mouse model of Kabuki Syndrome. Kabuki Syndrome results in ID and multisystem deficits including distinct craniofacial features and growth delays, but is not typically associated with ASD. It is caused by mutations in either KMT2D/MLL2, which methylates H3K4, or KDM6A, which demethylates H3K27 (Van Laarhoven et al., 2015). We examined RNA-seq data from a *Kdm6a* knockout mouse model, and, similar to Williams Syndrome models, found non-significant overlaps and enrichments by GSEA (**Fig. 5I-J**). These observations demonstrate that the transcriptional signature defined here is also disrupted in multiple mouse models of ASD, but is not observed more broadly in models of other neurodevelopmental disorders that only result in ID.

Next, we repeated these comparisons with the upregulated ASD gene signature. In this case, we found no significant overlap in any of the mouse models used here and only a single significant enrichment by log2 fold change pre-ranked GSEA (**Supplemental Fig. 7A-P**). These data suggest that the downregulated gene set as being a more relevant signature that is detected throughout multiple models of syndromic ASD. Finally, as an additional negative control, we repeated these analyses with a neurodegenerative disorder rather than only examining neurodevelopmental syndromes. We examined RNA-seq data from multiple mouse models of Huntington’s Disease which is caused by an expansion of the polyglutamine track in the Huntingtin (HTT) protein (Langfelder et al., 2016; Yildirim et al., 2019). We found either non-significant overlaps and GSEA enrichments, or in some cases, detected inverted enrichments where the downregulated gene signature overlapped and was enriched in genes upregulated in the mouse model (**Fig. 5K-L, Supplemental Fig. 8A-N).** Together, these data indicate that the downregulated transcriptional signature is disrupted in mouse models of NDDs that result in ASD, but not those that only result in ID, and is absent, or even reversed in neurodegenerative disorders. Conversely, the upregulated transcriptional signature detected in neuronal cultures is not found in animal models and thus may be more specific to downstream or compensatory changes that are unique to the cell culture experimental model used.

### Identification of the transcriptional signature in human ASD patient iPSCs

Given that the downregulated transcriptional signature was present in multiple animal models of ASD, we next asked whether this signature is expressed in human brain at times relevant to the development of ASD and whether these signature genes are also disrupted in human patients with ASD. To this end, we first used gene modules identified from BrainSpan data that are expressed at similar levels throughout the lifespan (Ji et al., 2016). We found that the downregulated transcriptional signature was highly enriched in gene modules 4, 6, and 36. These modules peak just before (36) or at (6) birth, or within the first months of life (4), all of which are time points highly relevant to the onset of ASD (**Supplemental Fig. 9A**). The upregulated transcriptional signature was enriched in modules 7 and 23 which are highly expressed in early development before birth (**Supplemental Fig. 9B**).

Next, we examined RNA-seq data from induced pluripotent stem cells (iPSCs) derived from idiopathic ASD or neurotypical patients, differentiated into neurons (DeRosa et al., 2018; Marchetto et al., 2017). Given the inherent variability in iPSCs obtained from unrelated individuals, very few significantly differentially expressed genes were detected in these datasets. We therefore used a recently developed RNA deconvolution method (Phan et al., 2020) to determine if a gene set of interest can distinguish disease state between multiple patient samples using principle component and linear regression analysis (Wright et al., 2017). Remarkably, we found that the downregulated transcriptional signature was sufficient to separate control and ASD iPSC-derived neurons based on expression of these signature genes (**Fig. 6A**). To confirm this finding wasn’t specific to the dataset used, we repeated this analysis with an additional available dataset from iPSC-derived neurons from control or ASD patients (DeRosa et al., 2018). While this dataset was insufficiently powered, similar trends were observed (**Fig. 6B**). In addition, we binned patient-derived neurons into two groups based on severity of ASD using ADOS score (Lord et al., 2000), which broadly assesses social interaction and communication abilities, and found that the downregulated transcriptional signature also was able to separate these two patient populations (**Supplemental Fig. 10A**). Conversely, but fitting with findings from animal models of ASD, the upregulated transcriptional signature was not able to distinguish between control and ASD human iPSC-derived neurons (**Supplemental Fig. 10B-D**). These data indicate that the downregulated transcriptional signature, defined within primary cultured neurons, was detected within human iPSC-derived neurons from idiopathic ASD patients. Together this work defines a transcriptional signature that encodes critical neuronal developmental proteins, contains unique chromatin features, and is present throughout multiple experimental models of ASD and in both mouse and human neurons.

**Figure 6.**
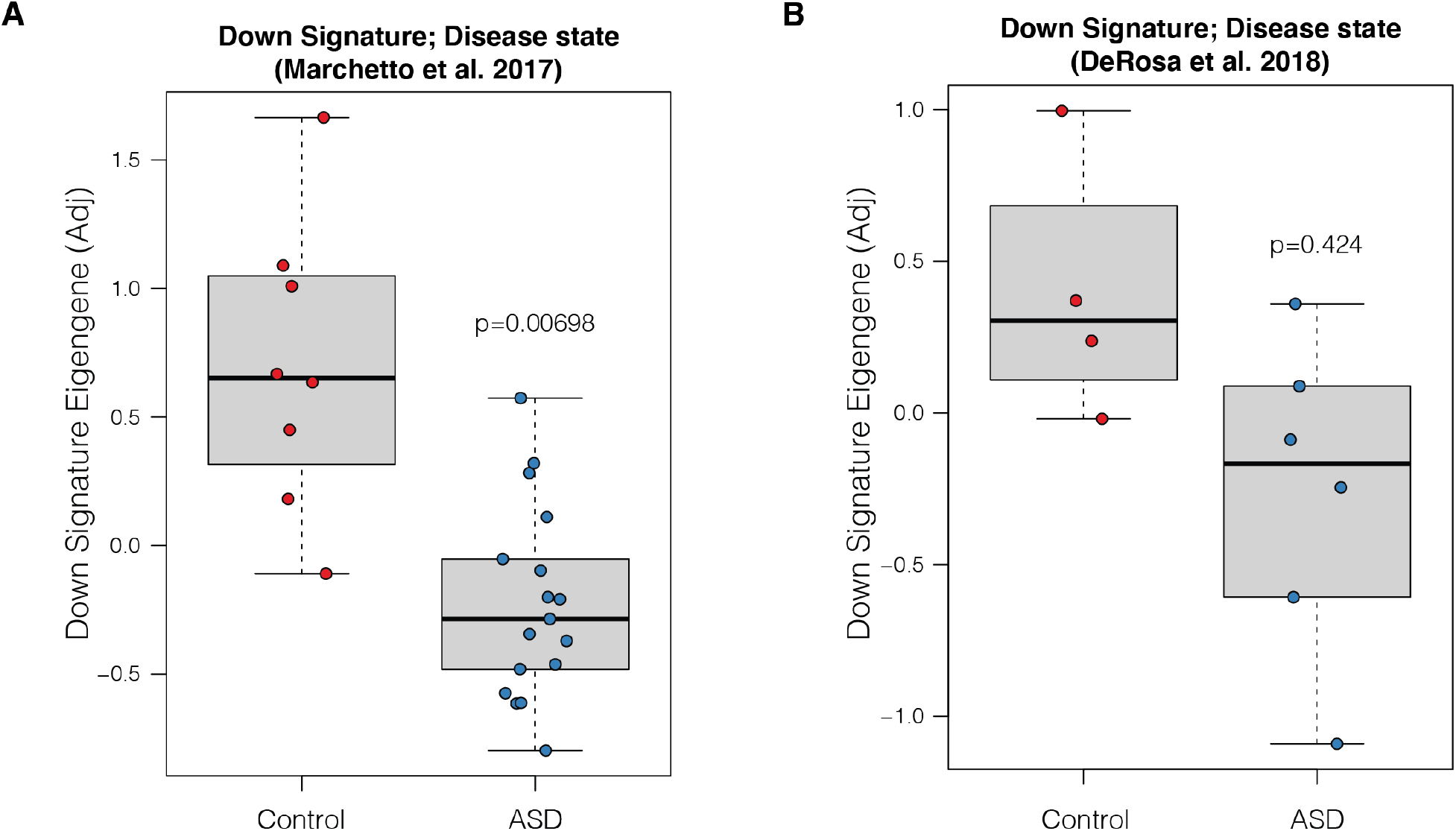
Identification of transcriptional signature in human iPSC-derived neurons with idiopathic ASD. (A) RNA deconvolution analysis of control and idiopathic ASD patient iPSC derived neurons (Marchetto et al. 2017) using the downregulated transcriptional signature (linear regression for disease state, p=0.00698). (B) RNA deconvolution analysis of control and idiopathic ASD patient iPSC derived neurons from an additional dataset (DeRosa et al. 2018) using the downregulated transcriptional signature (linear regression for disease state, p=0.424). Control indicates neurons derived from neurotypical human iPSCs. ‘Adj’ indicates adjusted.

## Discussion

Here we defined transcriptional signatures that are shared in response to knockdown of 5 chromatin associated proteins linked to neurodevelopmental disorders. Characterization of the function of these signatures demonstrated that these genes encoded proteins critical to neuronal development and function, with a notable enrichment in neurotransmitter transport genes and activity-dependent genes. In addition, the chromatin features associated with these signatures were enriched for specific histone modifications such as those encoding bivalent domains. Notably, the downregulated transcriptional signature was significantly enriched within several mouse models of ASD but not in mouse models of neurodevelopmental disorders that result in ID in the absence of ASD. Finally, the downregulated signature can distinguish between control and idiopathic ASD patient iPSC-derived neurons based on their transcriptional profiles.

A major finding that emerged from the analyses described here is that the *downregulated* transcriptional signature appears to be relevant to a wider range of models of ASD than the *upregulated* signature. We found that the downregulated signature genes map onto mouse models of neurodevelopmental disorders, particularly those linked to ASD, better than the upregulated signature genes in the context of cell culture models. The set of shared downregulated genes also was able to distinguish between control and ASD patient iPSC-derived neurons using an RNA deconvolution analysis, while upregulated genes were not, again suggesting that this downregulated gene set is the most relevant to ASD. Remarkably, these human iPSCs were generated from patients with idiopathic ASD, rather than from patients with a disorder resulting from loss of one of the 5 chromatin modifiers analyzed here. This indicates that this downregulated transcriptional signature is relevant to ASD more broadly and is not restricted to a subset of neurodevelopmental syndromes with defined genetic causes.

Interestingly, the downregulated transcriptional signature was detected within multiple models of syndromic ASD despite the highly divergent function of the chromatin modifiers chosen for analysis. This demonstrates that disruptions to chromatin that contribute to ASD result in similar gene expression changes regardless of the specific chromatin modifier disrupted and regardless of whether the disrupted histone modifications are associated with active or repressive gene expression. In addition, the 5 chromatin modifiers examined here are also putative targets of the Fragile X Mental Retardation Protein FMRP (Darnell et al., 2011), which has several functions, including regulation of translation of target transcripts. Interestingly, FMRP target transcripts are often upregulated in FXS when FMRP is lost (Korb et al., 2017), due to its role as a translational repressor. Conversely, in the neurodevelopmental syndromes modeled here, these targets typically have loss of function mutations or deletions. Despite these opposing mechanisms, the ASD signature we defined here maps onto both cellular and animal models of FXS. This suggests that regardless of the directionality of the disruptions to chromatin modifiers, many of the same gene expression changes are observed, and, even more unexpectedly, these genes are differentially expressed in the same direction. While each individual chromatin modifier and disease model also clearly had many additional genes unique to that model, this shared signature indicates that signature genes possess common features that underlie their sensitivity.

While the neuronal culture model used here allows for a highly controlled comparison of the effects of depletion of multiple chromatin modifiers in parallel, it also has several limitations. By its nature, it does not allow for the examination of how these chromatin modifiers may have distinct roles at different time points. It also does not take into account very early developmental defects that may result from mutations in chromatin modifiers even before cells differentiate into neurons. Further, this system focuses only on neurons in isolation and thus does not capture the more complex results of chromatin disruptions in multiple cell types in the brain and throughout the body. However, these limitations are also what allows for the direct and precise comparison of the immediate transcriptional effects of these chromatin modifiers without the confounding factors of system-wide disruptions and lifelong developmental deficits. In addition, the finding that the transcriptional signature identified in this system can be identified within the brain of multiple mouse models of related neurodevelopmental disorders suggests despite such limitations, the data described here still provide valuable insights into the link between chromatin misregulation, transcriptional disruptions, and ASD.

The features that cause the disruption of the transcriptional signature genes are not yet clear. However, the enrichment of specific chromatin states in these genes indicates that histone modifications may contribute to their sensitivity to the loss of chromatin modifying proteins. Similarly, past research has uncovered unusual chromatin features that are found both in the genes whose loss leads to ASD, as well as in genes whose expression is disrupted in idiopathic ASD (Zhao et al., 2018). In our analysis, we found several chromatin features of interest such as the presence of bivalent histone modifications and differences in modifications found at enhancers. Since many of the enzymes that control these modifications can be targeted by small molecule inhibitors, this finding raises the possibility of potentially targeting such modifications to reverse gene expression changes that ultimately lead to ASD. In summary, our data suggest the presence of a transcriptional signature found in multiple models of ASD and detected across multiple species. This indicates that common transcriptional disruptions may underlie the neuronal dysfunction that ultimately results in ASD and neurodevelopmental disorders.

## Materials and Methods

### Animals

All mice used were on the C57BL/6J background, housed in a 12 hr light/dark cycle, and fed a standard diet. All experiments were conducted in accordance with and approval of IUCAC.

### Primary Neuron Culture

Cortices were dissected from E16 C57BL/6J embryos and cultured in supplemented neurobasal medium (Neurobasal (Gibco 21103-049), B27 (Gibco 17504044), Glutamax (Gibco 35050-061), PenStrep (Gibco 15140-122)) in TC-treated 6-well plates coated with 0.05 mg/mL Poly-D-Lysine (Sigma Aldrich A-003-E).

### shRNA Knockdown

At 3 DIV, neurons were treated with 0.5 uM AraC. At 5 DIV, neurons were transduced overnight with lentivirus containing an shRNA sequence targeted to one of the ALCMs. Viral media was replaced the following day (6 DIV) and neurons were cultured for 5 additional days before downstream processing at 11 DIV.

### Lentivirus Production

HEK293T cells were cultured in high glucose DMEM growth medium (Corning 10-013-CV), 10% FBS (Sigma Aldrich F2442-500ML), and 1% PenStrep (Gibco 15140-122)). Calcium phosphate transfection was performed with Pax2 and VSVG packaging plasmids. shRNAs in a pLKO.1-puro backbone were purchased from Sigma Aldrich Mission shRNA library (SHCLNG). Viral media was removed 12 hours post-transfection and collected at 24 and 48 hours later. Viral media was passed through a 0.45 μm filter and precipitated overnight with Pegit solution (40% PEG-8000 (Sigma Aldrich P2139-1KG), 1.2 M NaCl (Fisher Chemical S271-1)). Viral particles were pelleted and resuspended in 200μL PBS.

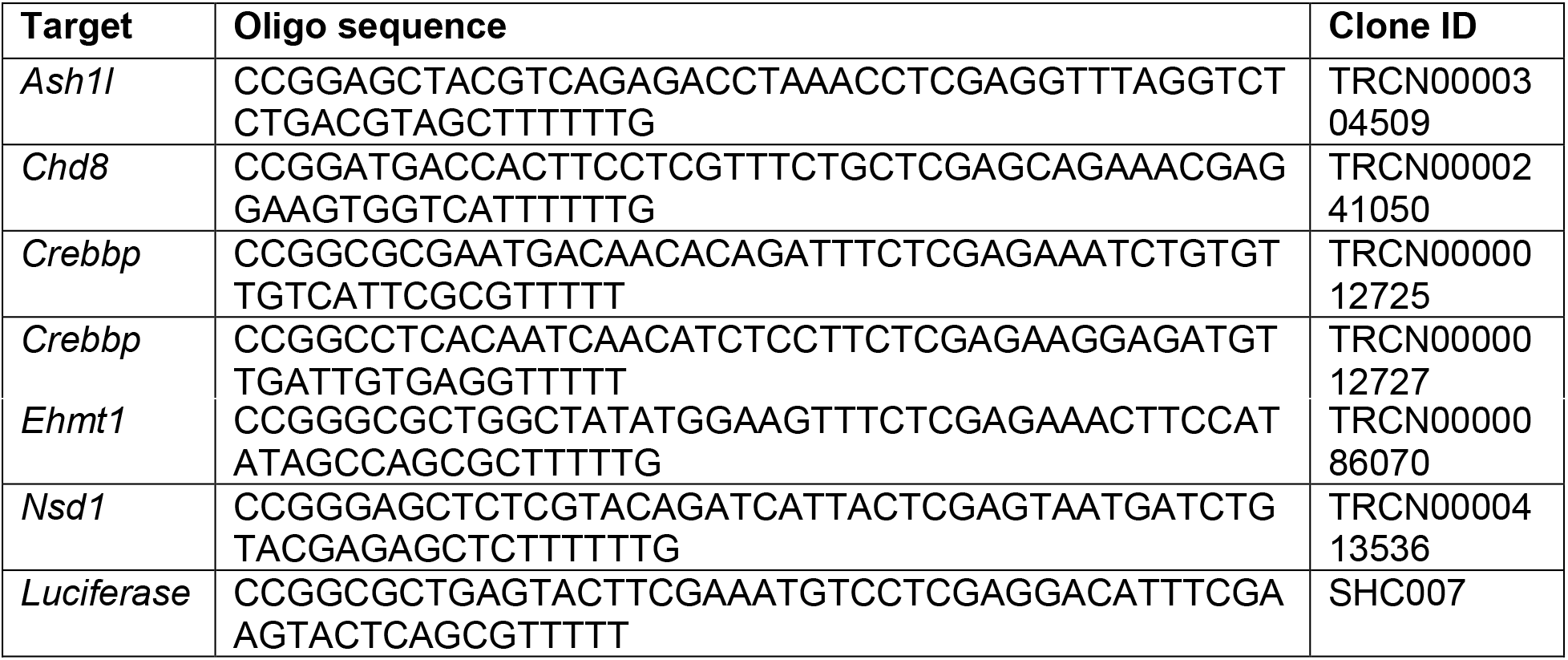

### RNA isolation

Total RNA was collected from each transduction at 11 DIV for both RT-qPCR and RNA-seq. RNA for RT-qPCR was isolated from neurons using Qiagen RNeasy mini kit (74004) or Zymo Quick-RNA miniprep kit (R1054).

### Western blotting

After 10 DIV, neurons were lysed in RIPA (25 mM Tris pH 7.6, 150 mM NaCl, 1% NP-40, 1% sodium deoxycholate, 0.1% SDS), supplemented by protease inhibitor (Roche 04693124001), phosphatase inhibitor (Roche 04906837001), 1mM DTT, 1mM PMSF. Lysates were mixed with 5X Loading Buffer (5% SDS, 0.3 M Tris pH 6.8, 1.1 mM Bromophenol blue, 37.5% glycerol), boiled for 10 minutes, sonicated for 5 minutes, and cooled on ice. Sample protein was resolved by 16% Tris-Glycine, 4-20% Tris-Glycine, or 3-8% Tris-Acetate SDS-PAGE, followed by transfer to a 0.45μm PVDF membrane (Sigma Aldrich IPVH00010) for immunoblotting. Membranes were blocked for 1 hour in 5% BSA in TBST and probed with primary antibody overnight at 4C.

| Target | Supplier (Cat No.) | Working dilution |
| --- | --- | --- |
| ASH1L | Bethyl Laboratories (A301-749A) | 1:1000 |
| CHD8 | Cell Signaling Technologies (77694S) | 1:1000 |
| CREBBP | Cell Signaling Technologies (7389S) | 1:1000 |
| EHMT1 | Thermo Fisher Scientific (PA5114733) | 1:1000 |
| TUBULIN | Abcam (ab18207) | 1:5000 |
| HRP Goat anti-Rabbit | Abcam (ab6721) | 1:10000 |
| HRP Sheep anti-Mouse | Millipore Sigma (GENA931-1ML) | 1:10000 |

### RT-qPCR

cDNA was prepared with High-Capacity cDNA Reverse Transcription Kit (Applied Biosystems 4368813) and quantitative PCR was performed with Power SYBR Green PCR Master Mix (Applied Biosystems 4367659).

Data was analyzed using the Common Base Method developed by Ganger et al. 2017. Reported statistics were calculated by student’s T-test (two-tailed, heteroscedastic) based on primer efficiency-weighted deltaCT values (ACLM-vs. Luciferase-infected). Reported bar graph values represent the square root of the relative expression ratio for a given gene of interest.

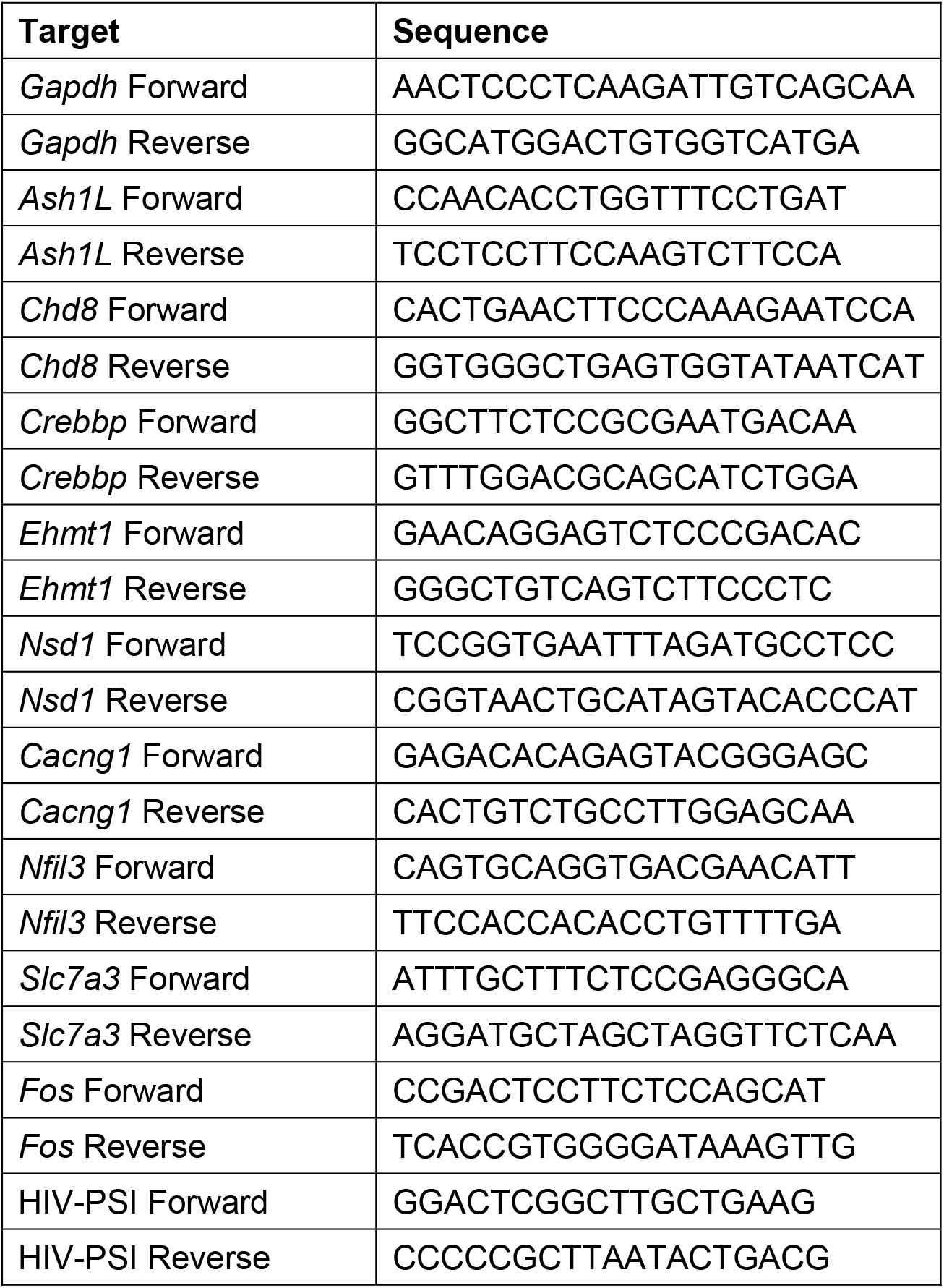

### RNA-seq

Libraries were generated using the Illumina TruSeq Stranded mRNA Library Prep Kit (Illumina 20040534). Libraries were sequenced on an Illumina NextSeq 500/550 and reads (75-basepair single-end) were mapped to Mus Musculus genome build mm10 with Salmon, The R package, DESeq2 (v3.14), was used to perform differential gene expression analysis. IGV tools (2.11.3) was used to generate genome browser views.

### Statistical analyses

#### Overlap analysis

R was used to calculate pairwise overlaps of differentially expressed gene lists and their corresponding significance values. For these analyses, a background expressed gene list was defined to be those genes in the Luciferase versus Uninfected DESeq2 comparison with an official gene symbol and a baseMean value greater than 5. InveractiveVenn was used to generate 5-way gene list Venn diagrams (Heberle et al., 2015). Multiple testing correction based on a 5% False Discovery Rate threshold using the Benjimini-Hochberg method (Benjamini and Hochberg, 1995).

#### Gene Ontology analysis

HOMER (v4.11) was used to perform gene ontology analysis using default parameters. GeneWalk (v1.5.3) was run using Python 3.9 using default values. According default cutoffs, GeneWalk results were filtered for global_padj and gene_padj values less than 0.1 (Ietswaart et al., 2021). Revigo was used to remove redundant terms and to generate a 2-dimensional scatter based on semantic similarity of retained terms. Revigo input parameters used were: size of resulting list – tiny; remove obsolete GO terms – yes; species – mus musculus; semantic similarity measure – Resnik. Python 3.9 was used for processing of Revigo outputs with semantic similarity scatters clustered iteratively using Kmeans (sklearn.cluster.KMeans; default parameters; 2-15 clusters). Resulting clusters were assessed on the basis of silhouette scores and by manually judging the biological similarity of ontology terms placed within the same clusters.

#### ChromHMM analysis

The chromatin state segmentation BED file containing coordinates of annotated chromatin states (Gorkin et al., 2020) was used in running ChromHMM with parameters: java -mx400M ChromHMM OverlapEnrichment inputsegment inputcoorddir outfileprefix. Input coordinates for mm10 genic regions were generated via the UCSC Table Browser (group: Genes and gene predictions; track: GENCODE VM11 (Ensembl 86); table: basic). Mouse transcript names were converted to corresponding mouse Ensembl IDs using the R package biomaRt. Genic coordinates were obtained by querying the a biomaRt mart object (ensembl) for ‘start_position’ and ‘end_position’ and filtering for genes of interest. Promoter coordinates were generated using the R package TxDb.Mmusculus.UCSC.mm10.knownGene (‘knownGene’). The flank function from the GenomicRanges R package was used to generate coordinates 500 and 2000 base pairs upstream of TSS sites listed in knownGene. The cds function from GenomicRanges was used to get cds coordinates from knownGene. Promoter regions at −500 and −2000 base pairs were trimmed if they intersected a genic region.

#### GSEA

The R package fgsea was used to perform pre-ranked gene set enrichment analysis based on log2FoldChanges obtained from DESeq2 differential expression analysis. The ASD Down and ASD Up gene lists were used as comparison gene sets.

#### RNA deconvolution

RNA deconvolution analysis was performed as described in Phan et al. 2020. Code was obtained from the Lieber Institute Github page (https://github.com/LieberInstitute/PTHS_mouse). Variance Stabilized Transformation (DESeq2 package) was used to scale gene expression values which were used to compare read counts between ASD and control samples. Linear regression analysis was performed on transformed gene expression values, modeling the effect of diagnosis while adjusting for age and a number of quality surrogate variables (qSVs), which help correct for additional confounds in the data and are determined by the num.sv() function in the sva R package (Jaffe et al., 2017). Given limited access to metadata, we performed this analysis after removing variation due to age of the patient. The optimal number of PCs for Marchetto et al., 2017 diagnosis and ASD-severity comparisons were determined to be 7 and 4, respectfully. For DeRosa et al. 2018, the subset of induced-neurons cultured for 35 DIV was used. The optimal number of PCs for diagnosis comparisons was determined to be 4.

## Data availability

All genome-wide sequencing data is available under accession number GSE193663.

---

**Supplemental Figure 1.**
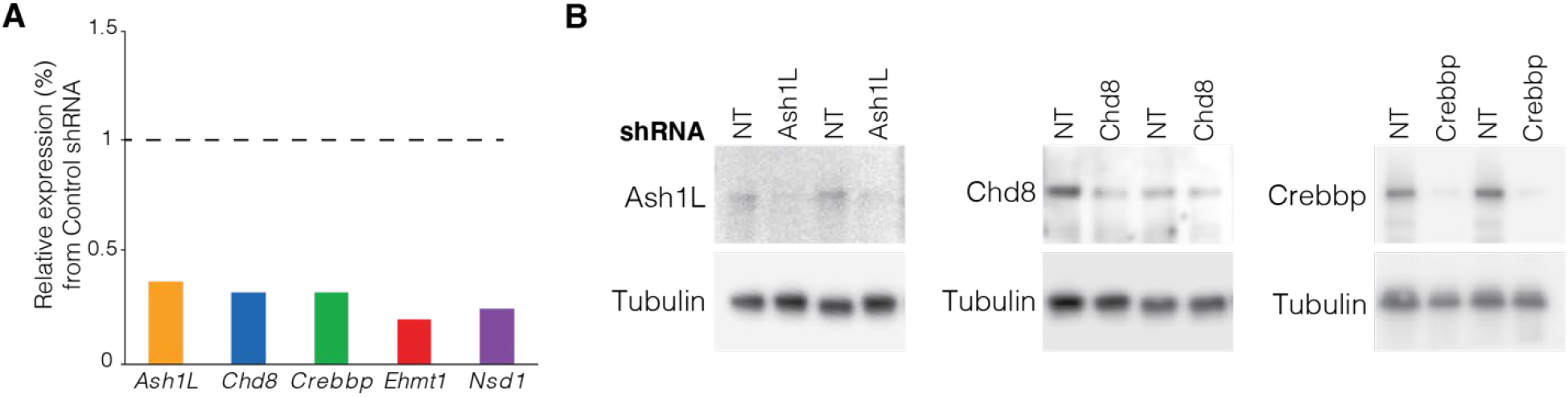
Confirmation of knockdown of target chromatin modifiers. (A) RT-qPCR analysis of knockdown after infection with shRNA lentiviral vectors. N = 3. (B) Western blot analysis of protein levels of ASD-linked chromatin modifiers targeted by shRNA lentiviruses. NT indicates non-targeting control lentivirus infection.

**Supplemental Figure 2.**
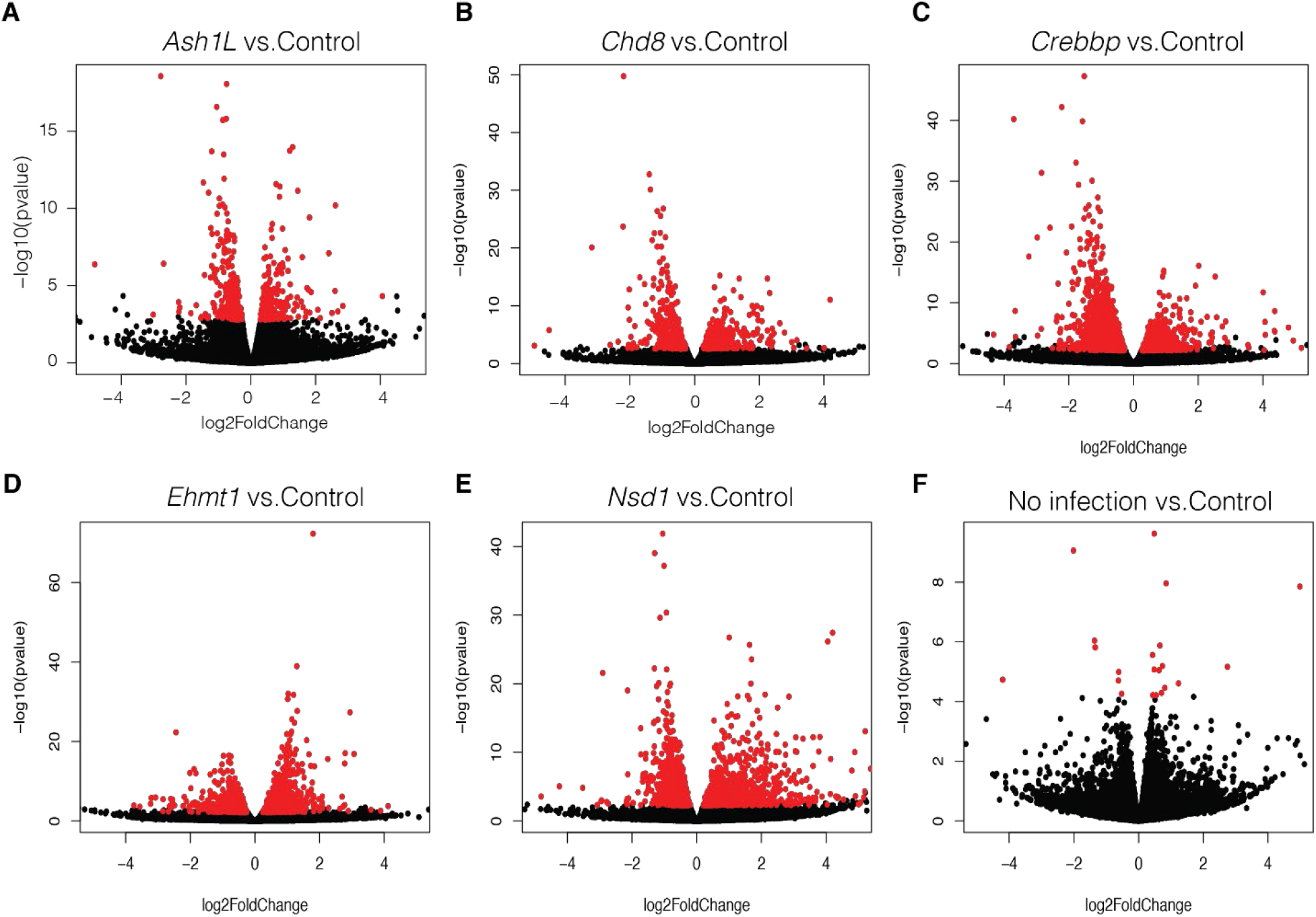
Differential gene expression in response to knockdown of ASD-linked chromatin modifiers. Volcano plot of differential gene expression after knockdown of 5 ASD-linked chromatin-associated proteins by infection of shRNA lentivirus compared to non-targeting control lentivirus. N=3. Red indicates significance at an adjusted p-value of 0.05 by DESeq2.

**Supplemental Figure 3.**
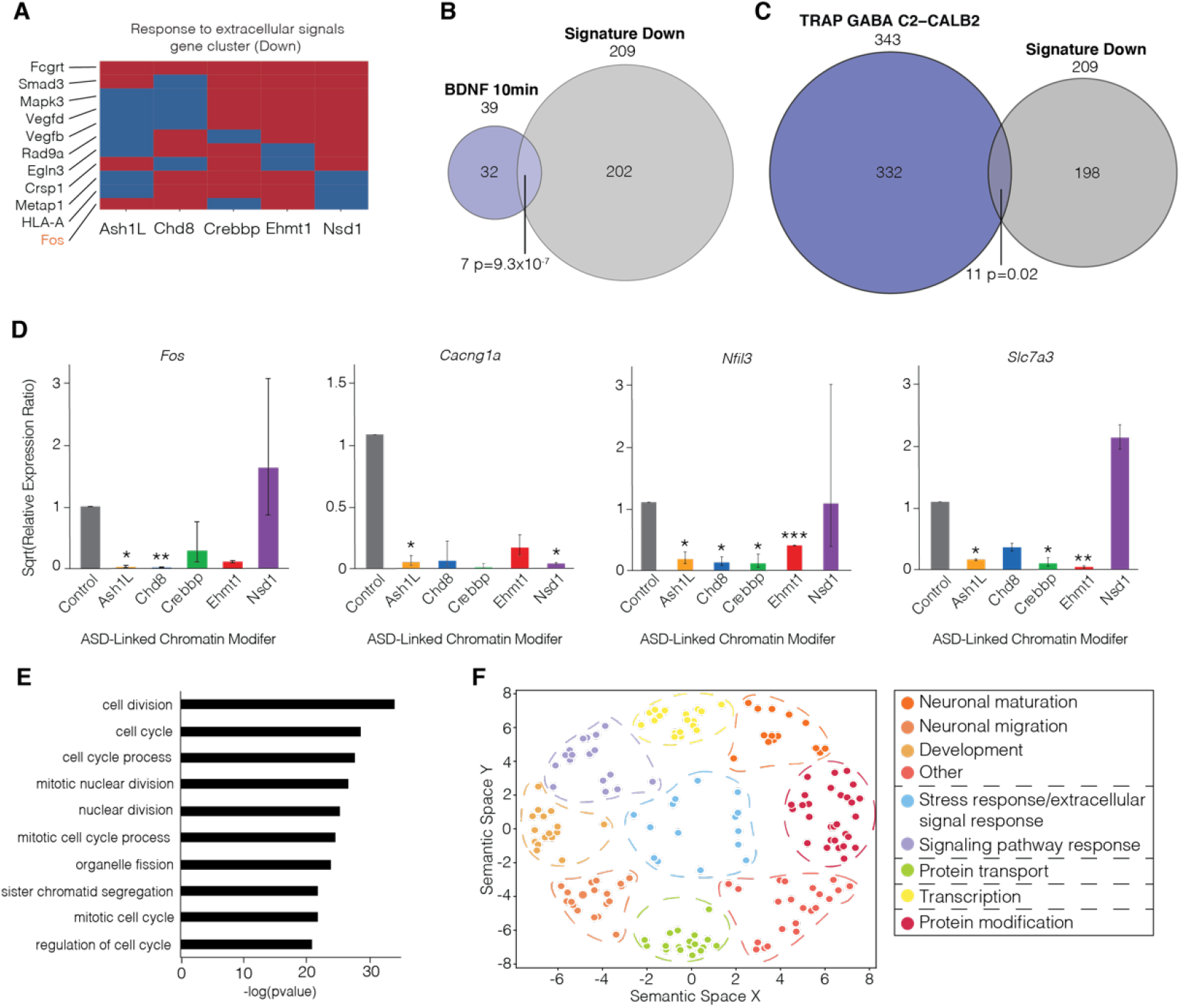
Function of down and upregulated transcriptional signature genes. (A) Genes contributing to ‘Response to extracellular signals’ cluster that are differentially expressed after knockdown of 3 or more ASD-linked chromatin modifiers. Activity dependent genes shown in orange. (B) Overlap of downregulated transcriptional signature genes with genes induced in response to a 10-minute BDNF stimulation in primary cultured neurons. (C) Overlap of downregulated transcriptional signature genes with genes induced in a fear conditioning memory paradigm in neurons activated in a TRAP2 mouse model. (D) RT-qPCR validation of genes that are disrupted by at least 3 of the 5 ASD-linked chromatin modifiers and that contribute to gene ontology clusters. N=3. (E) Gene Ontology analysis of upregulated transcriptional signature gene function. (F) GeneWalk analysis followed by Revigo clustering of upregulated transcriptional signature genes. Overlap significance determined by hypergeometric tests. RT-qPCR statistics determined by t-test of means of CT values normalized to *Gapdh* relative to control infection, * indicates <0.05, ** indicates <0.01, *** indicates <0.001. BDNF indicates Brain Derived Neurotrophic Factor.

**Supplemental Figure 4.**
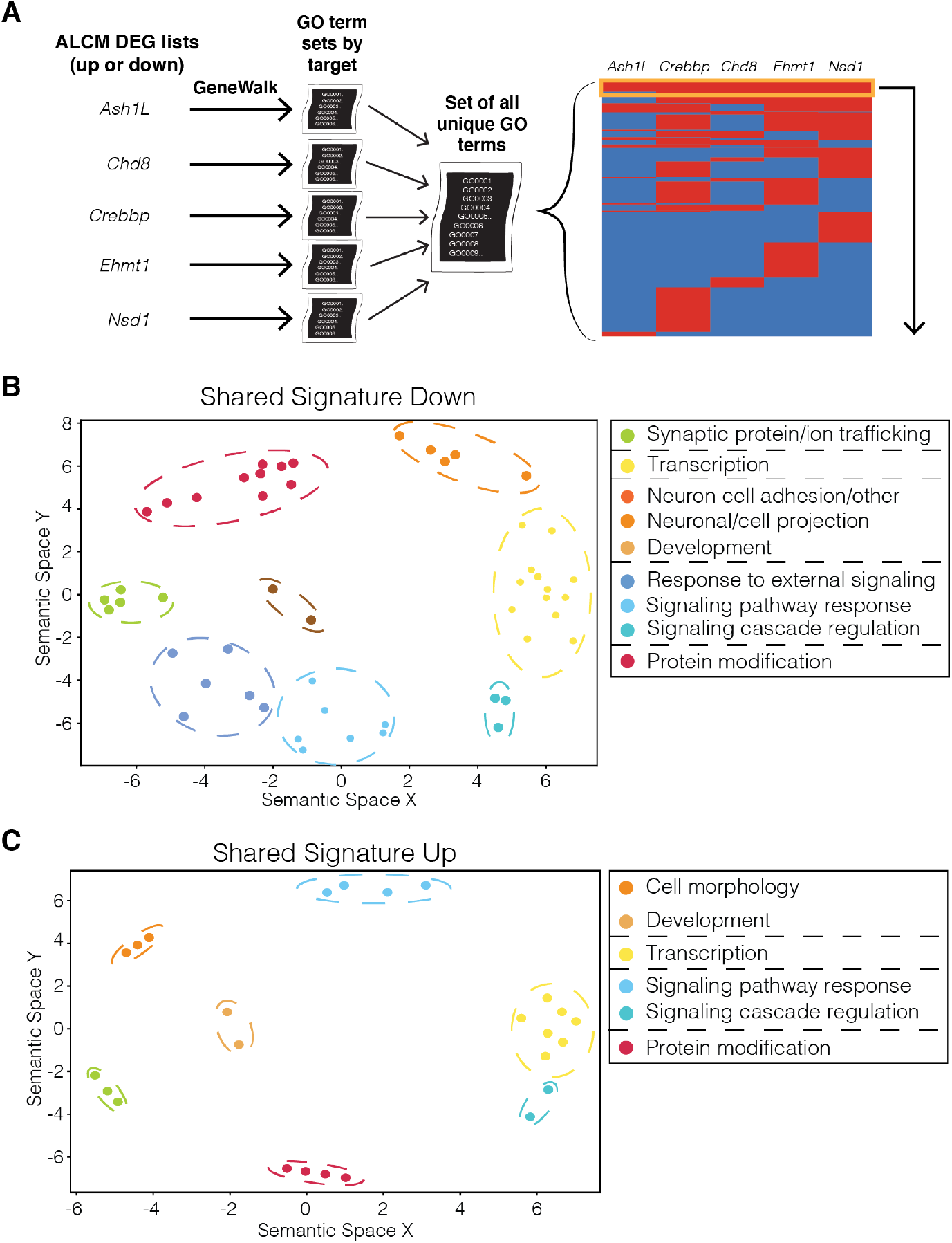
Function of down and upregulated genes for each ASD-linked chromatin modifier. (A) Analysis schematic of GeneWalk performed on each separate set of differentially expressed genes following knockdown of 5 ASD-linked chromatin modifiers (ALCM). GO terms were then overlapped to find common functions and clustered by REVIGO. (B) Analysis of separate GeneWalk analysis and overlapping outputs of genes downregulated following knockdown of ASD-linked chromatin modifiers. (C) Analysis of separate GeneWalk analysis and overlapping outputs of genes upregulated following knockdown of ASD-linked chromatin modifiers. DEG indicates differentially expressed genes following knockdown of an ALCM target compared to non-targeting control lentiviral infection.

**Supplemental Figure 5.**
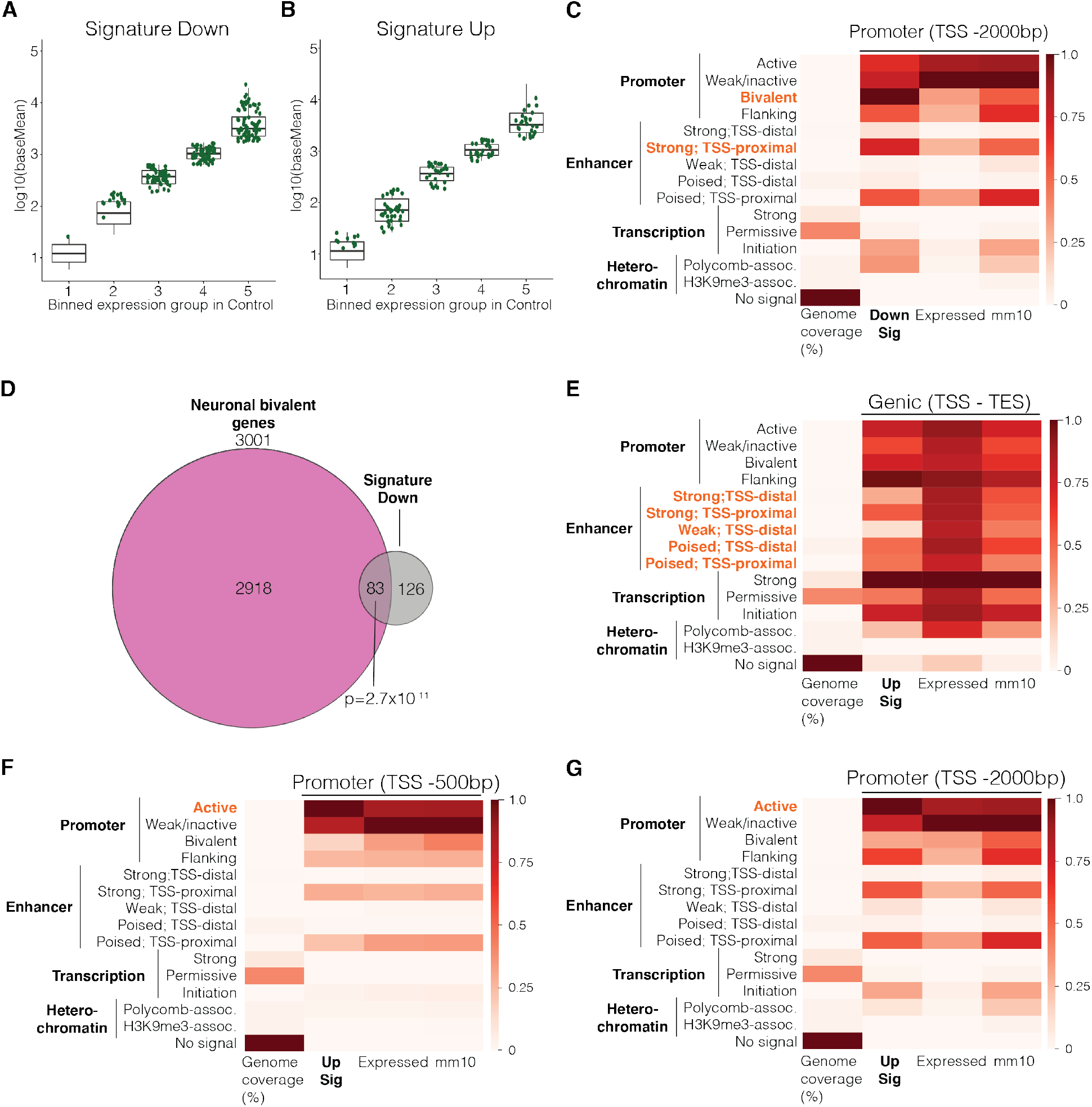
Chromatin states in transcriptional signature genes. (A-B) Expression of all genes from control neurons binned into 5 equal groups with down (A) and up (B) ASD transcription signature genes shown in green. (C) Downregulated transcriptional signature genes overlap with bivalent genes expressed in neurons. Overlap significance based on a hypergeometric test. (D) ChromHMM analysis of promoter region of downregulated transcriptional signature genes using an expanded upstream region up to 2000 basepairs upstream of the TSS. (E) ChromHMM analysis of genic regions of upregulated transcriptional signature genes. (F) ChromHMM analysis of the promoter region of upregulated transcriptional signature genes up to 500 base pairs upstream of the TSS. (G) ChromHMM analysis of the promoter region of upregulated transcriptional signature genes using an expanded upstream region up to 2000 base pairs upstream of the TSS. TSS indicates transcription start site. TES indicates transcription end site. Expressed indicates genes expressed in neuronal culture system. Displayed heatmaps represent overlap enrichment output values range-normalized by column.

**Supplemental Figure 6.**
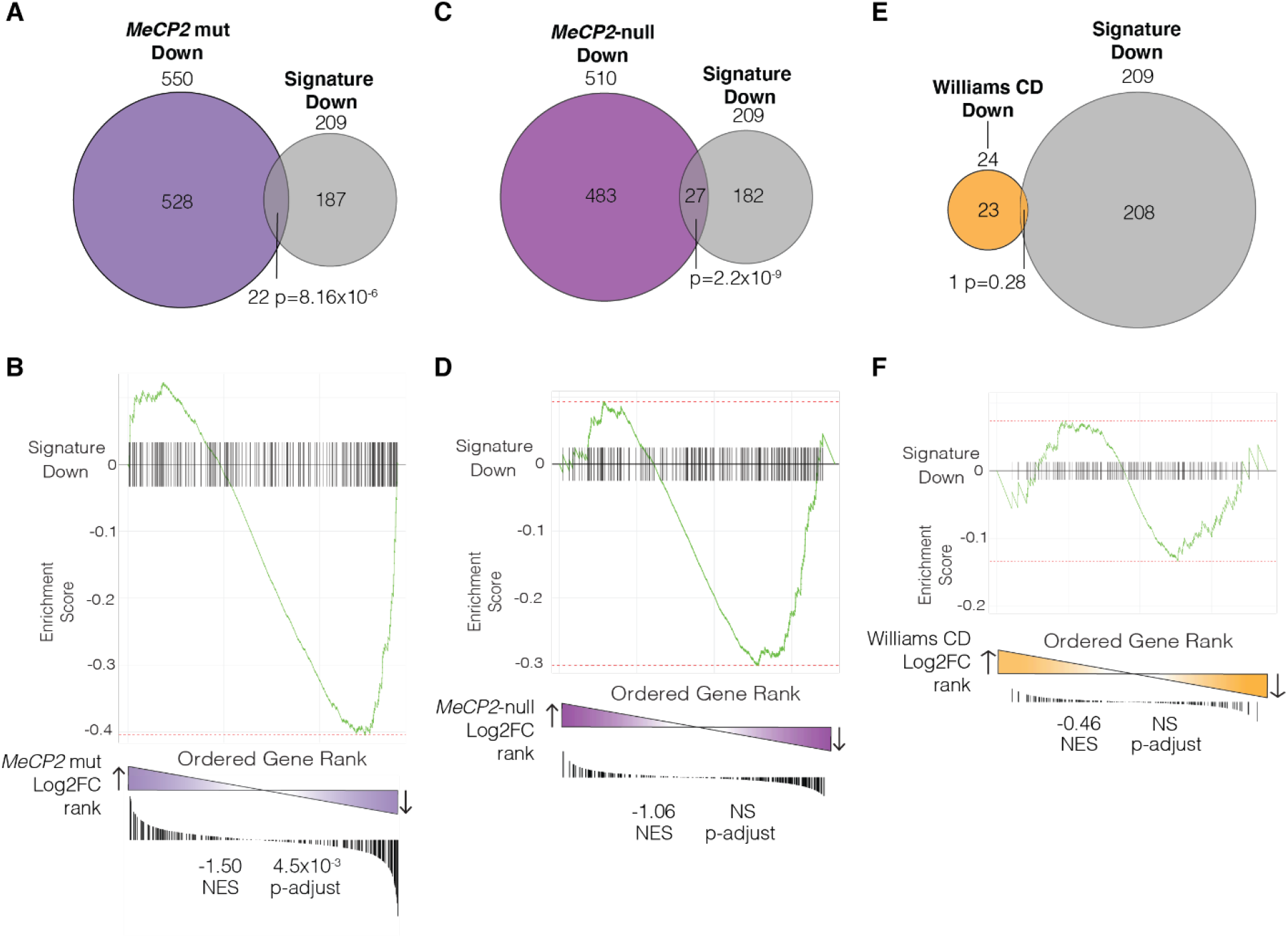
Examination of downregulated transcriptional signature in additional mouse models of NDDs. (A-B) Overlap (A) and GSEA (B) analysis of downregulated transcriptional signature compared to differentially expressed genes in a mouse model of Rett Syndrome containing a mutated *MeCP2* gene (T158M), frequently seen in human RTT patients. (C-D) Overlap (C) and GSEA (D) analysis of downregulated transcriptional signature compared to differentially expressed genes in a mouse model of Williams Syndrome containing the full deletion comparable to that seen in human patients. Overlap significance based on hypergeometric tests. ‘mut’ indicates *MeCP2* T158M mutation commonly found in cases of RTT. CD indicates complete deletion on 5G2 analogous to the human Williams Syndrome Critical Region on 7q11.23. NES indicates normalized enrichment score.

**Supplemental Figure 7.**
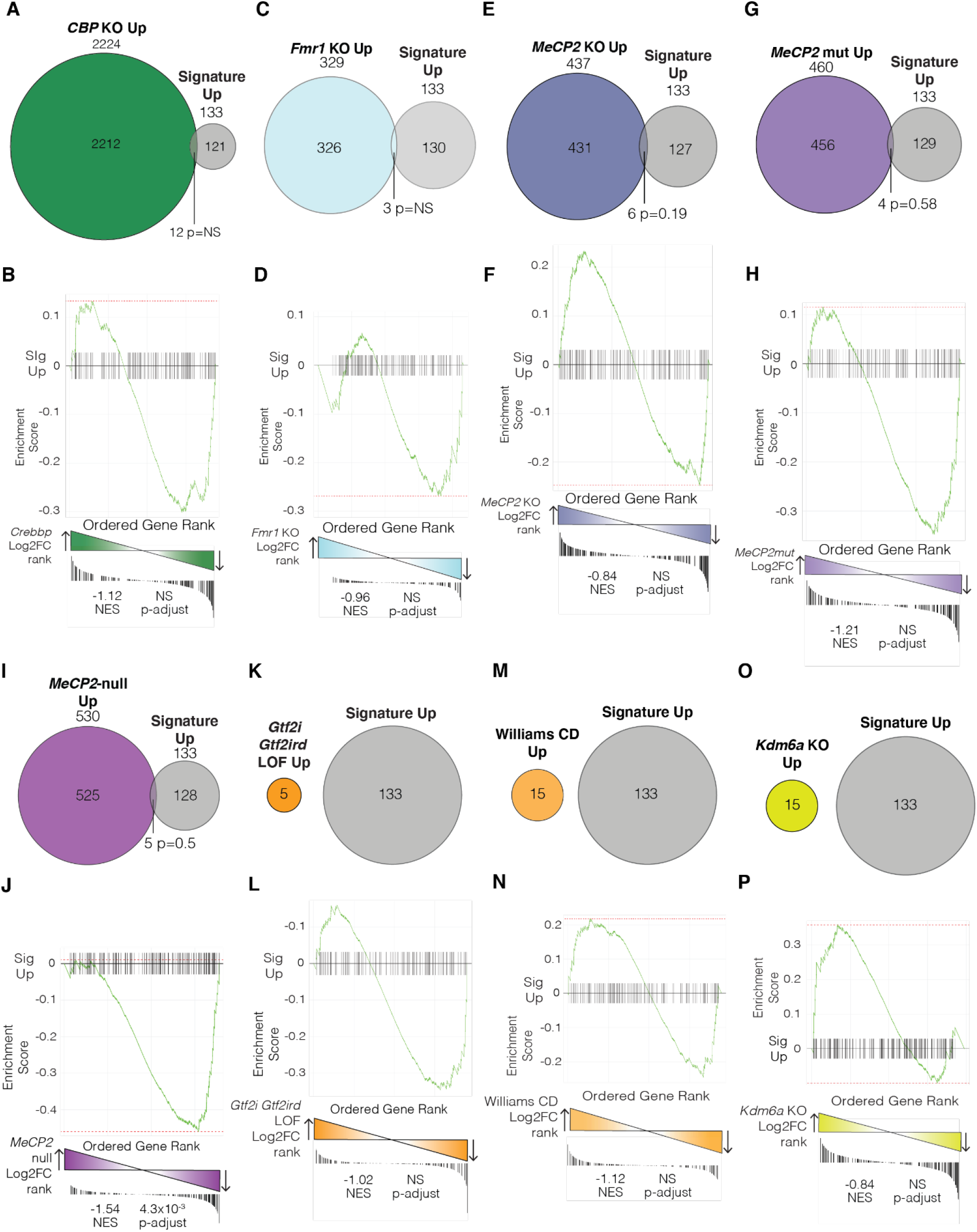
Examination of upregulated transcriptional signature in mouse models of NDDs. (A-B) Overlap (A) and GSEA (B) analysis of upregulated transcriptional signature compared to differentially expressed genes in a *Kat3a* double KO mouse model of Rubinstein-Taybi Syndrome. (C-D) Overlap (C) and GSEA (D) analysis of upregulated transcriptional signature compared to differentially expressed genes in a *Fmr1* KO mouse model of FXS. (E-F). Overlap (E) and GSEA (F) analysis of upregulated transcriptional signature compared to differentially expressed genes in a *MeCP2* KO mouse model of Rett Syndrome. (G-H) Overlap (G) and GSEA (H) analysis of upregulated transcriptional signature compared to differentially expressed genes in a mouse model of Rett Syndrome containing a mutated *MeCP2* gene (T158M). (I-J) Overlap (I) and GSEA (J) analysis of upregulated transcriptional signature compared to differentially expressed genes in a *Gtf2i* and *Gtf2ird* double LOF mouse model of Williams Syndrome. (K-L) Overlap (K) and GSEA (L) analysis of upregulated transcriptional signature compared to differentially expressed genes in a mouse model of Williams Syndrome containing the full deletion comparable to that seen in human patients. (M-N) Overlap (M) and GSEA (N) analysis of upregulated transcriptional signature compared to differentially expressed genes in a *Kdm6a KO* mouse model of Kabuki Syndrome. Overlap significance based on hypergeometric tests. CD indicates complete deletion on 5G2 analogous to the human Williams Syndrome Critical Region on 7q11.23. ‘mut’ indicates *MeCP2* T158M mutation commonly found in cases of RTT. NES indicates normalized enrichment score.

**Supplemental Figure 8.**
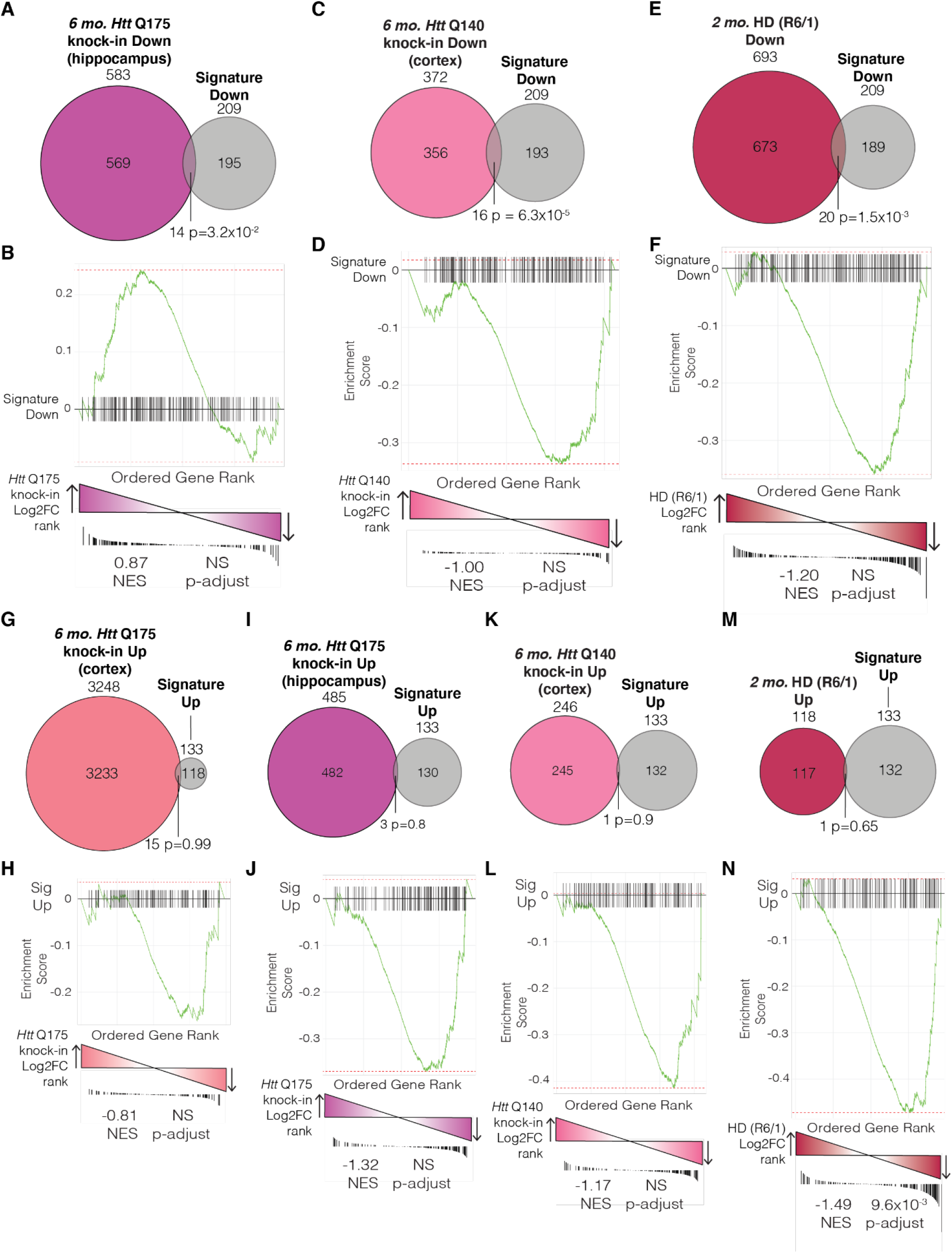
Examination of transcriptional signature in mouse models of Huntington’s Disease. (A-B) Overlap (A) and GSEA (B) analysis of downregulated transcriptional signature compared to differentially expressed genes from the hippocampus of a mouse model of Huntington’s Disease containing 175 glutamine repeats. Mice were aged 6 months along with littermate WT controls. (C-D) Overlap (C) and GSEA (D) analysis of downregulated transcriptional signature compared to differentially expressed genes from the cortex of a mouse model of Huntington’s Disease containing 140 glutamine repeats. Mice were aged 6 months along with littermate WT controls. (E-F) Overlap (E) and GSEA (F) analysis of downregulated transcriptional signature compared to differentially expressed genes from the R6/1 mouse model of Huntington’s Disease containing 115 glutamine repeats. Mice were aged 2 months along with age-matched controls. (G-H) Overlap (G) and GSEA (H) analysis of upregulated transcriptional signature compared to differentially expressed genes from the cortex of a mouse model of Huntington’s Disease containing 175 repeats (corresponding to Fig. 5K-L) (I-J) Overlap (I) and GSEA (J) analysis of upregulated transcriptional signature compared to differentially expressed genes from the hippocampus of a mouse model of Huntington’s Disease containing 175 repeats. (K-L) Overlap (K) and GSEA (L) analysis of upregulated transcriptional signature compared to differentially expressed genes from the cortex of a mouse model of Huntington’s Disease containing 140 repeats. (M-N) Overlap (M) and GSEA (N) analysis of upregulated transcriptional signature compared to differentially expressed genes from the R6/1 mouse model of Huntington’s Disease containing 115 repeats. Overlap significance based on hypergeometric tests. NES indicates normalized enrichment score.

**Supplemental Figure 9.**
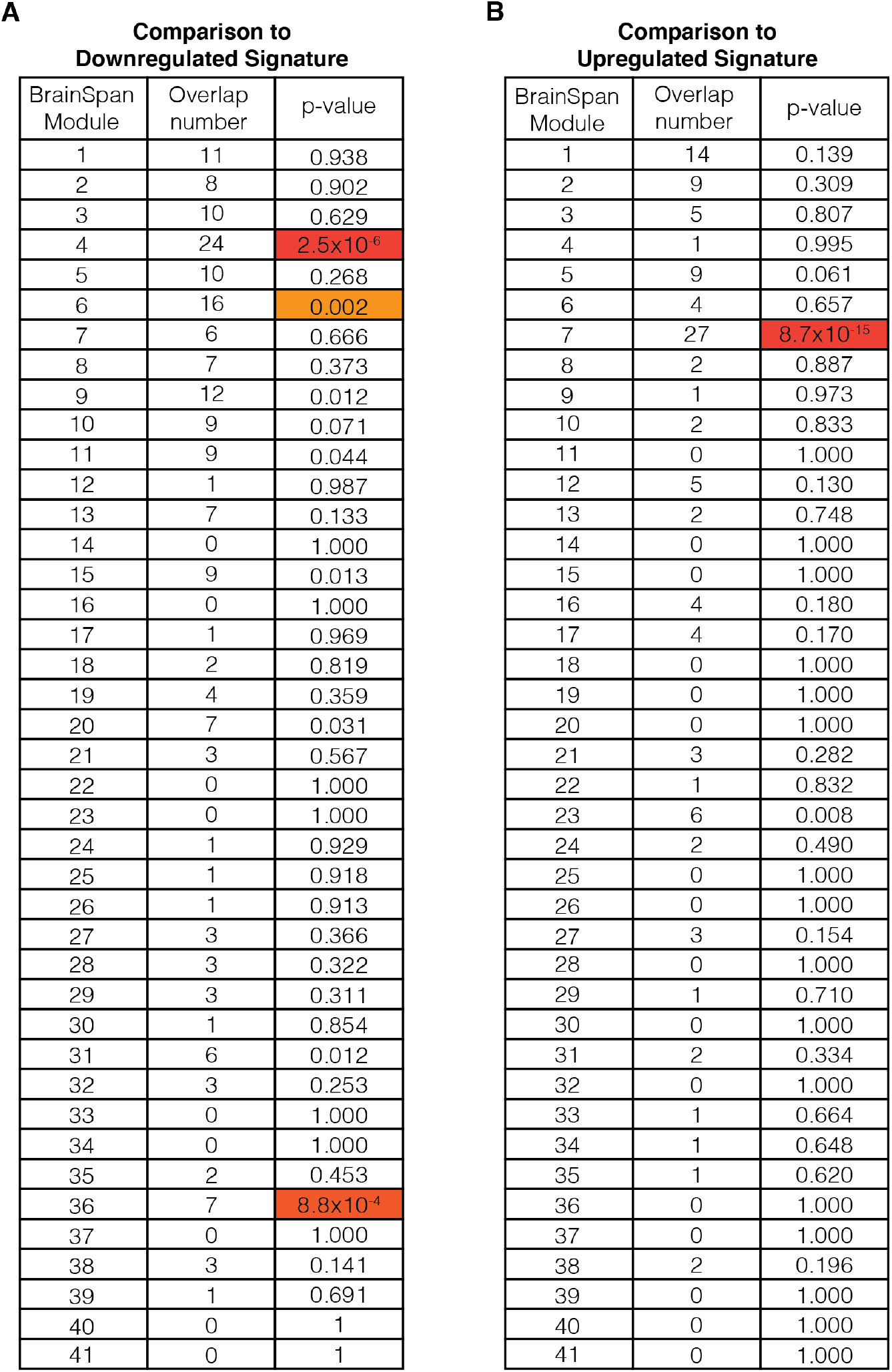
Developmental expression of transcriptional signature. (A-B) Overlap of the downregulated (A) or upregulated (B) transcriptional signature with modules of co-expressing genes identified from human BrainSpan data. Overlap significance based on hypergeometric tests with significant overlaps highlighted in orange. Significance based on 5% FDR threshold corrected for multiple comparisons using the Benjamini-Hochberg method.

**Supplemental Figure 10.**
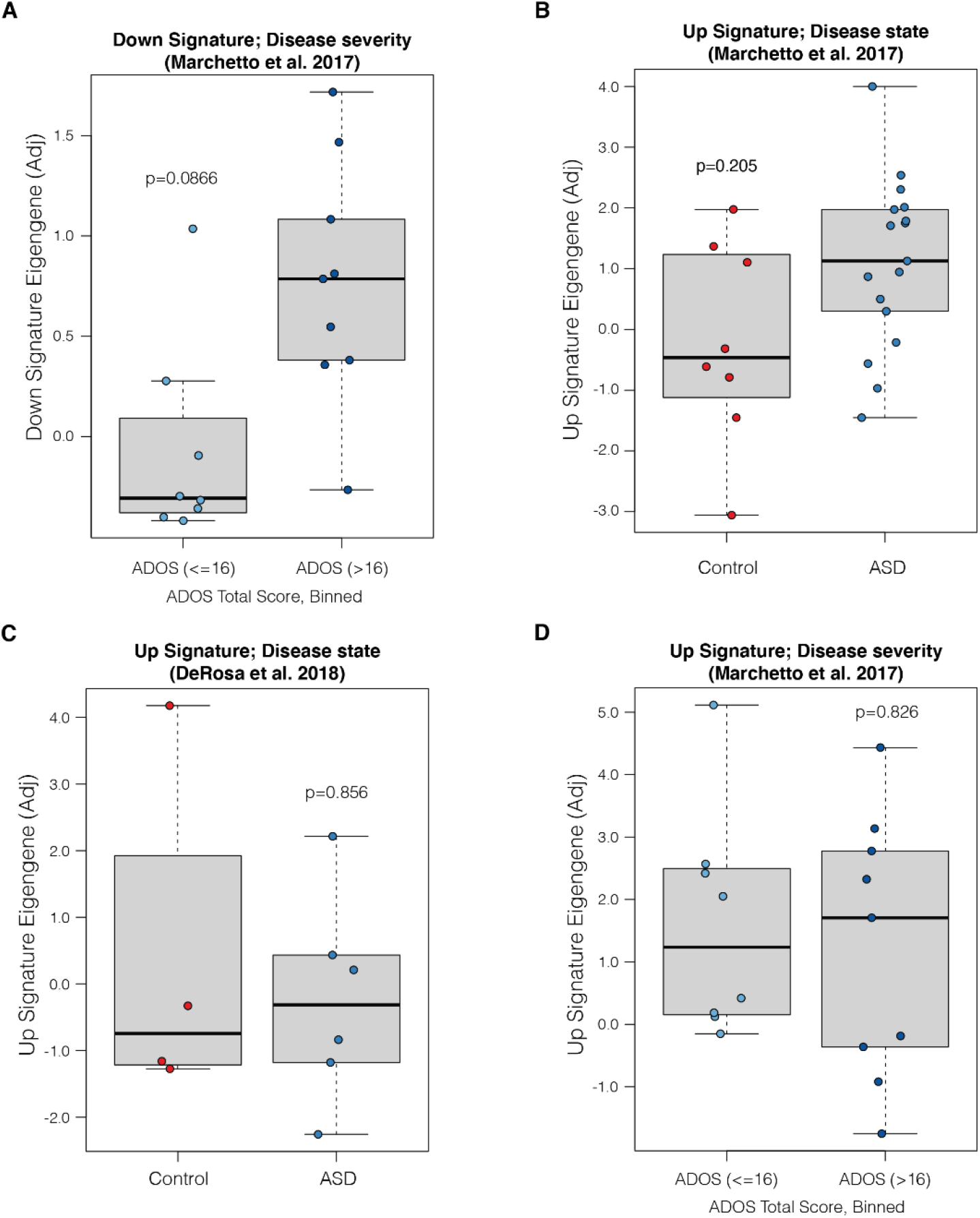
Transcriptional signature in human iPSC-derived neurons with idiopathic ASD. (A) RNA deconvolution analysis of idiopathic ASD patient iPSC derived neurons separated by ADOS score using the downregulated transcriptional signature. (B) RNA deconvolution analysis of control and idiopathic ASD patient iPSC derived neurons (Marchetto et al., 2017) using the upregulated transcriptional signature. (C) RNA deconvolution analysis of control and idiopathic ASD patient iPSC derived neurons from an additional dataset (DeRosa et al., 2018) using the upregulated transcriptional signature. (D) RNA deconvolution analysis of idiopathic ASD patient iPSC derived neurons separated by ADOS score using the upregulated transcriptional signature. Control indicates neurons derived from neurotypical human iPSCs. ‘Adj’ indicates adjusted.

